# Long lived liver-resident memory T cells of biased specificities for abundant sporozoite antigens drive malaria protection by radiation-attenuated sporozoite vaccination

**DOI:** 10.1101/2024.11.08.622604

**Authors:** Maria N. de Menezes, Zhengyu Ge, Anton Cozijnsen, Stephanie Gras, Patrick Bertolino, Irina Caminschi, Mireille H. Lahoud, Katsuyuki Yui, Geoffrey I. McFadden, Lynette Beattie, William R. Heath, Daniel Fernandez-Ruiz

## Abstract

Vaccination with radiation-attenuated sporozoites (RAS) can provide highly effective protection against malaria in both humans and mice. To extend understanding of malaria immunity and inform the development of future vaccines, we studied the protective response elicited by this vaccine in C57BL/6 mice. We reveal that successive doses of *Plasmodium berghei* RAS favour the generation of liver CD8^+^ tissue-resident memory T cells (T_RM_ cells) over circulating memory cells, and markedly enhance their longevity. Importantly, RAS immunisation strongly skews the composition of the liver CD8^+^ T_RM_ compartment towards cells specific for abundant sporozoite antigens, such as thrombospondin-related adhesive protein (TRAP) and circumsporozoite protein (CSP), which become major mediators of protection. The increased prevalence of sporozoite specificities is associated with limited intrahepatic parasite development and inhibition of naïve T cell responses to all parasite antigens in previously vaccinated mice. This leads to the exclusive expansion of effector T cells formed upon initial immunisation, ultimately reducing the diversity of the liver T_RM_ pool later established. These findings provide novel insights into the mechanisms governing malaria immunity induced by attenuated sporozoite vaccination and highlight the susceptibility of this vaccine to limitations imposed by strain-specific immunity associated with the abundant, yet highly variable sporozoite antigens CSP and TRAP.

**Author Summary:** Malaria remains a significant global health challenge. An efficient vaccine could significantly enhance malaria control. Vaccination with radiation-attenuated sporozoites (RAS) can induce highly efficient protection against malaria, and our study brings critical insights into the protective mechanisms elicited by this vaccine. We show that RAS stimulates the formation of parasite-specific cytotoxic memory T cells that permanently reside in the liver (liver T_RM_ cells). These cells are critical mediators of protection. Interestingly, multiple doses of RAS extend the lifespan of these memory cells, potentially improving long term immunity. However, we found that the induced memory T cell response is strongly skewed towards abundant, but highly variable, sporozoite proteins. Thus, this phenomenon exposes a potential limitation of the RAS vaccine against the great parasite diversity in the field, as it focuses the T cell response away from less abundant, but more conserved, parasite antigens.

## Introduction

Malaria still kills over 600,000 individuals annually, with 75% of these fatalities occurring among children under five years of age in low income countries [1]. Vaccines stand out as one of the most efficient and cost-effective public health interventions against infectious diseases [2], particularly in resource-limited regions. Radiation attenuated sporozoites (RAS) is one of the most effective malaria vaccines, demonstrating long lived sterilising protection across mice, non-human primates (NHP), and humans against *Plasmodium* spp. sporozoite infection [3-5]. This vaccination approach involves the inoculation of sporozoites attenuated through exposure to either X- or gamma-irradiation. Attenuation is achieved by a carefully calibrated sublethal radiation dose that enables parasites to invade the liver after injection into the host but induces their subsequent arrest in this organ, inhibiting progress to the blood stage [6-8].

Considerable research has been dedicated to elucidating the protective mechanisms that underlie RAS-induced protection. RAS vaccination requires multiple doses for maximal efficacy [9], and elicits both humoral and cellular immunity. Although the former can contribute to protection [10-13], cellular immunity, mediated by CD8^+^ T cells, is a pivotal component of RAS-mediated immunity, as demonstrated by the susceptibility to infection of different strains of vaccinated mice, and NHP, when these cells are removed [14-16]. RAS vaccination of mice and NHP was found to induce the accumulation of memory CD8^+^ T cells in the liver [13, 17-19]. Memory T cells can be subdivided into two major subtypes: circulating memory cells (T_CIRCM_), in turn separated into central (T_CM_) and effector (T_EM_) memory T cells depending on their recirculating pattern, and tissue-resident memory T cells (T_RM_), which remain in the organ wherein they are formed [20]. Although both types of memory CD8^+^ T cells are elicited by RAS vaccination and can contribute to protection [21, 22], tissue-resident memory T cells appear to be critical in C57BL/6 mice [22], which are highly susceptible to infection by *P. berghei* and require higher doses of RAS for sterilising immunity than other strains [23, 24]. This notion is also supported in humans, as sterilising protection is still detected in repeatedly vaccinated individuals despite their antibody levels or numbers of circulating memory CD8^+^ T cells being comparable to those in their unprotected counterparts [13, 19, 25].

CD8^+^ T_RM_ cells must find and kill parasites during the brief period of residence in the liver, which lasts 2 days in mice and 7 days in humans [26-28]. An essential determinant of the capacity of CD8^+^ T_RM_ cells to exert anti-parasitic protection is their antigen specificity, as parasite antigens must be presented via Major Histocompatibility Complex class I (MHC-I) molecules on hepatocytes to enable CD8^+^ T_RM_ cell recognition of infected cells. *Plasmodium* protein expression vastly changes throughout different stages of the parasite’s life cycle [29, 30], and T_RM_ cells specific for antigens with contrasting expression patterns therefore differ in their protective capacity. Thus, T_RM_ cells that recognise the 60S ribosomal protein L6 (RPL6), an antigen predominantly expressed during liver stage, exhibit heightened protective efficacy relative to those specific for thrombospondin-related adhesive protein (TRAP) [31], which is mainly expressed by sporozoites [30, 32]. A broad diversity of T cell specificities is thought to promote better protection [33], as late antigens with prolonged expression patterns provide a longer window of opportunity for T cells to find and eliminate parasites before they progress to the blood. Other requirements defining an antigen’s immunogenicity and protective capacity are its efficient processing by the proteasome for the generation of peptides that can be loaded onto MHC-I molecules, the strong and stable binding of these peptides to MHC molecules, and the presence of naïve T cells in the endogenous repertoire capable of responding to the peptide/MHC-I complexes generated [34, 35]. Notably, C57BL/6 mice are not known to mount CD8^+^ T cell responses specific for *P. berghei* circumsporozoite protein (CSP) [36], another major sporozoite antigen that is also expressed during liver stage.

Dissecting the dynamics of the protective T cell responses elicited by RAS is crucial for understanding immunity to liver stage malaria. In this study, we sought to investigate the landscape of memory T cell specificities triggered by successive *P. berghei* RAS vaccinations in C57BL/6 mice. Our findings reveal a progressive skewing of the liver T_RM_ response towards sporozoite antigen specificities, which undergo robust expansion and become major mediators of protection against live sporozoite challenge. Surprisingly, this immunodominance is established even for T cells targeting the sporozoite antigen TRAP, despite the modest intrinsic protective capacity of these cells. Additionally, repeated immunisations significantly extend the half-life of the parasite-specific T_RM_ cells generated while suppressing naïve T cell responses to any parasite antigen. These elements limit the breadth of the ensuing memory response by largely constraining the induced liver T_RM_ cell pool to cells specific for previously encountered sporozoite antigens.

## Results

### Liver CD8^+^ T_RM_ cells protect RAS-vaccinated C57BL/6 mice against *P. berghei* sporozoite infection

Using MHC-I restricted, TCR transgenic PbT-I cells [37], specific for the *Plasmodium* antigen RPL6 [31], we previously showed that the protection provided by two doses of RAS, administered 30 days apart, is largely mediated by a liver-resident subset of memory CD8^+^ T cells [22]. However, the vaccination schedule utilised was suboptimal, only providing ∼40% of sterilising protection, and the presence of adoptively transferred PbT-I cells might have altered the endogenous protective responses naturally elicited by this vaccine [38]. Thus, to further characterise the mechanisms responsible for malaria immunity induced by RAS, we injected mice with smaller doses of irradiated parasites (10,000 sporozoites) at shorter intervals (one week apart), as this immunisation strategy is known to provide highly efficient sterilising protection against *P. berghei* sporozoite infection in C57BL/6 mice [39]. Mice were injected with 1, 2 or 3 weekly doses of 10,000 *P. berghei* RAS, and challenged with 200 live, fully infectious *P. berghei* sporozoites 30 days after the last RAS injection. A single injection of RAS (1xRAS) failed to induce sterilising protection against sporozoite infection, but mice in this group displayed significantly reduced parasitemia compared to unvaccinated controls on day 7 after challenge (Figure 1A, B). Protection was improved in mice vaccinated with two doses of RAS (2xRAS), which showed significant sterile protection and further reduced day 7 parasitemia. Three RAS injections (3xRAS) conferred even higher sterilising protection against sporozoite challenge (Figure 1A, B). Importantly, in line with previous findings in C57BL/6 mice and non-human primates [14, 16], depletion of CD8^+^ T cells in 3xRAS vaccinated mice completely removed protection (Figure S1A-C). Furthermore, parasitemia on day 7 in depleted mice was similar to that in unvaccinated controls, underscoring the major role of CD8^+^ T cells in protection in this system (Figure S1A-C). Next, we sought to define the contribution of liver T_RM_ cells to protection. These cells were selectively depleted in 3xRAS vaccinated mice via treatment with monoclonal antibodies specific for the T_RM_ surface marker CXCR3 [22] prior to challenge with *P. berghei* sporozoites. This treatment efficiently removed endogenous CD8^+^ T_RM_ cells in the liver, but left numbers of effector memory CD8^+^ T cells unaltered (Figure S1D-H). Numbers of central memory T cells were moderately reduced in the spleen, as some of these cells also express CXCR3, but not in the liver. As previously observed [22], parasite-specific T_CM_ cells are minimally induced by RAS vaccination in this model and are hence unlikely to play a significant role in protection (Figure S1F-H). Vaccinated mice treated with αCXCR3 mAb and depleted of liver T_RM_ cells became fully susceptible to sporozoite challenge, their parasitemias approaching those of unvaccinated mice (Figure 1C, D). This result strongly indicated that protection in this system is largely CD8^+^ liver T_RM_-dependent. Overall, these experiments confirmed that multiple RAS vaccinations can confer highly efficient protection against *Plasmodium* sporozoite infection in C57BL/6 mice and provided strong evidence that this protection is mediated by CD8^+^ T_RM_ cells.

**Figure 1.**
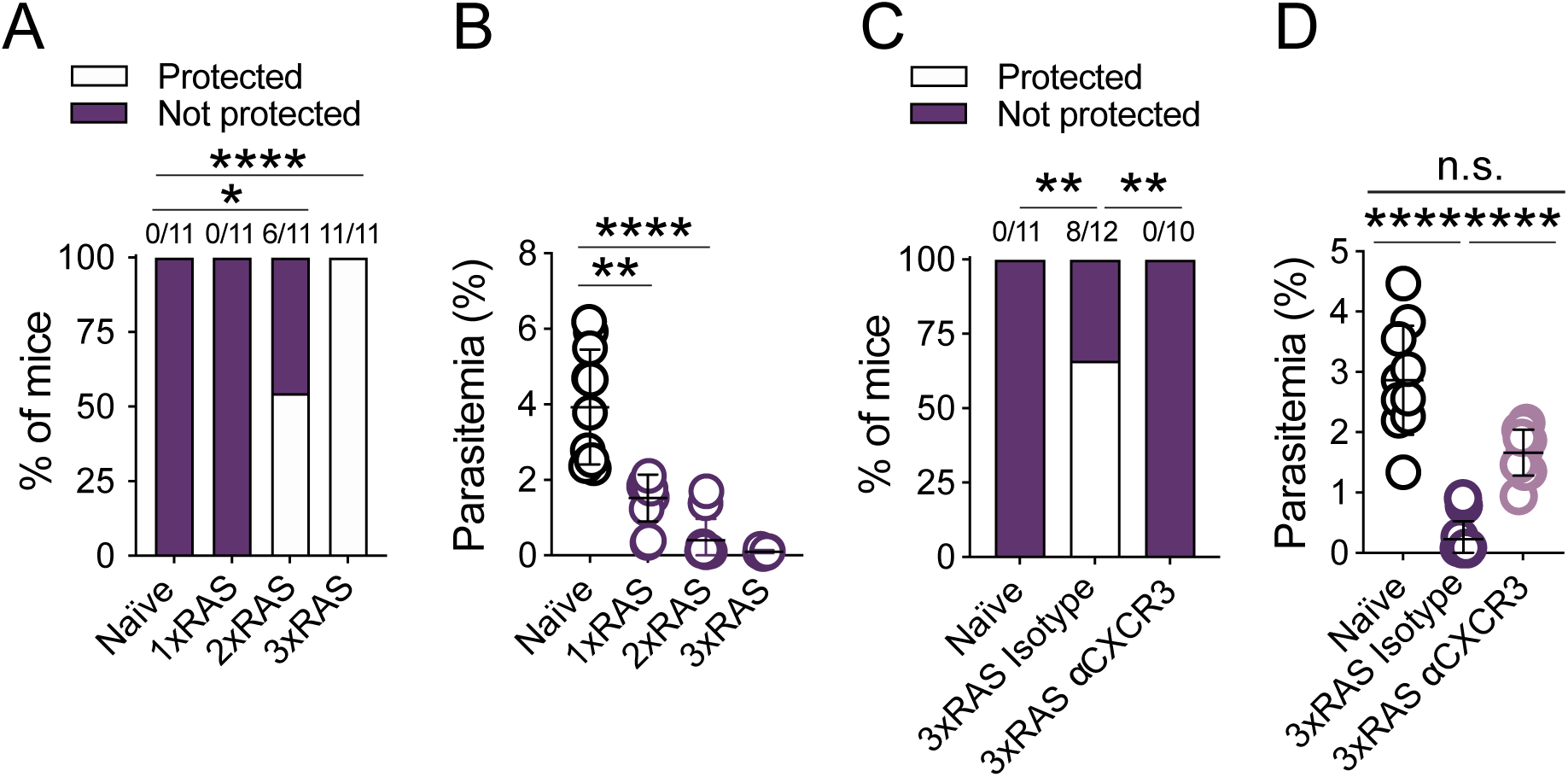
Repeated RAS vaccination of C57BL/6 mice provides efficient protection against *P. berghei* sporozoite infection that is dependent on CD8^+^ T_RM_ cells. (**A-B)** Mice were vaccinated with 1, 2 or 3 doses of 10,000 RAS (1xRAS, 2xRAS and 3xRAS respectively), one week apart and were challenged with 200 *P. berghei* sporozoites 30 days after the last dose of RAS. (**A**) Rates of sterile protection. Numbers above columns denote numbers of protected mice / total numbers of mice per group (**B**) Parasitemia at day 7 post-challenge. (**C-D**) Mice were vaccinated using three doses of 5,142-10,000 RAS, one week apart, and treated with αCXCR3 mAb 3 and 1 days prior to challenge with 200 live sporozoites, which was performed on day 30-34 after the last vaccination. Parasitemia was monitored to evaluate protection. (**C)** Rates of sterilising protection. (**D)** Parasitemia on day 7 after sporozoite infection. Data were pooled from 2 independent experiments. Comparisons of sterile protection rates were done using Fisher’s exact tests. Parasitemia data were log-transformed and compared using one-way ANOVA and Tukey’s multiple comparisons test.

### RAS boosting favours generation of liver CD8^+^ T_RM_ cells specific for abundant sporozoite antigens

As a whole parasite vaccine, RAS stimulates CD8^+^ T cell responses of multiple specificities [18, 33]. To better understand the features of the protective CD8^+^ T cell response elicited by repeated RAS vaccination, we next sought to determine the relative abundance of endogenous memory T cells specific for known parasite antigens generated in 3xRAS vaccinated mice. We focused on antigens with contrasting expression patterns during parasite development in the mouse; i) Thrombospondin-related anonymous protein (TRAP, PBANKA_1349800), containing the PbTRAP_130-138_ epitope [40], which is expressed at high levels by sporozoites [32, 41]; ii) the putative 60S ribosomal protein L6 (RPL6, PBANKA_1351900), an antigen predominantly expressed during liver and blood stage, but less abundantly in sporozoites, that contains the cognate antigen of PbT-I cells, PbRPL6_120-127_ [31, 37]; and iii) the replication protein A1 (RPA1, PBANKA_0416600), originally identified as a blood stage T cell antigen [42] but also expressed during liver stage (Figure S2A), that contains the PbRPA1_199-206_ epitope, also known as F4 [42]. Intriguingly, the size and composition of the memory CD8^+^ T cell compartment generated was markedly different depending on the number of RAS doses administered. Thus, one dose of 10,000 RAS generated similar numbers of RPL6-specific and TRAP-specific liver T_RM_ cells (Figure 2A, S2B), as well as circulating cells of these specificities in the liver and the spleen (Figure S2B). RPA1-specific cells were also expanded, indicating that this antigen is immunogenic during pre-erythrocytic stages, and this was also the case for total numbers of memory CD8^+^ T cells of undefined specificities (Figure 2A-C, S2B). However, a second injection of RAS profoundly changed the relative abundance of memory T cell specificities. TRAP-specific cells, particularly TRAP T_RM_ cells in the liver, were significantly boosted, with an average 23-fold increase vs a single RAS injection (Figure 2A-C, S2B). In comparison, RPL6-specific cells were only increased 3.6-fold, and RPA1-specific cells, or those of undefined specificities, remained largely unchanged (Figure 2C). This trend became more pronounced in mice injected with three doses of RAS (Figure 2A-C, S2B). Numbers of TRAP-specific T_RM_ cells were 77-fold higher in 3xRAS vaccinated mice compared to 1xRAS (Figure 2C) and constituted on average more than 30% of total liver T_RM_ cells (Figure 2B). In contrast, RPL6 and RPA1 specific cells expanded 7.5 to 3.5-fold vs 1xRAS (Figure 2C) and only accounted for 4.5% and 2% of all liver T_RM_ cells respectively (Figure 2B). For all specificities, repeated injections of RAS progressively favoured the generation of T_RM_ cells in the liver over circulating memory cells (T_CIRCM_, comprising T_EM_ and T_CM_ cells), as exemplified by the increasing ratios of liver T_RM_ vs spleen T_CIRCM_ cell numbers upon successive rounds of RAS vaccination (Figure 2D). These results showed that repeated vaccination with RAS favoured the generation of liver-resident memory CD8^+^ T cells specific for an abundant sporozoite antigen (i.e. TRAP) over those responding to antigens more prominently expressed at later stages.

**Figure 2.**
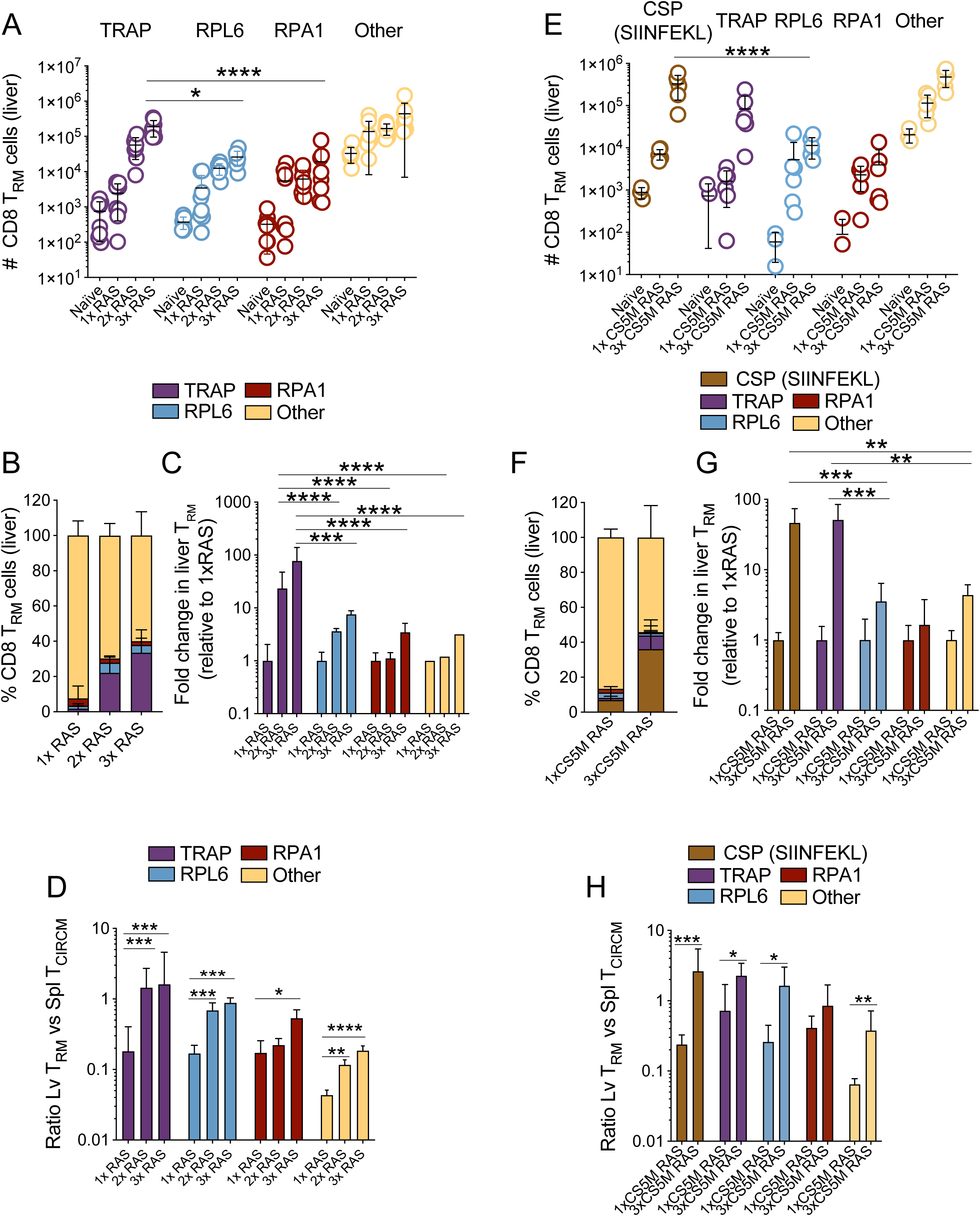
Abundance of liver CD8^+^ T_RM_ cells specific for known *Plasmodium* antigens in mice vaccinated multiple times with WT RAS. **A-D.** CD8^+^ T_RM_ cell responses specific for TRAP, RPL6, RPA1, or other specificities (tetramer-negative T_RM_ cells) in the livers of mice vaccinated with 1, 2 or 3 rounds of 10,000 RAS, one week apart. Cell numbers were assessed 30 days after the last round of immunisation. Data were log-transformed and compared using two-way ANOVA and Tukey’s multiple comparisons test. **A.** Liver T_RM_ cell numbers. Data were log transformed and statistically compared using two-way ANOVA and Tukey’s multiple comparisons test. **B.** Frequencies of liver T_RM_ cells of different specificities amongst all T_RM_ cells in the liver. **C.** Fold change in the number of T_RM_ cells of the indicated specificities compared to those in 1xRAS vaccinated mice. Data were log transformed and statistically compared using one-way ANOVA and Tukey’s multiple comparisons test. Data in A-C were pooled from two independent experiments. **D.** Ratios of T_RM_ cell numbers in the liver vs numbers of circulating memory T cells (T_CIRCM_, calculated by adding T_CM_ and T_EM_ cells) in the spleen. Data were compared using one-way ANOVA and Tukey’s multiple comparisons test. **E-H.** CD8^+^ T_RM_ cells specific for mutated CSP (SIINFEKL), TRAP, RPL6, RPA1, or other specificities in the livers of mice vaccinated with 1 or 3 rounds of 5,050-7,700 CS5M RAS, 4-8 days apart. Cell numbers were assessed 30-63 days after the last round of immunisation. **E.** Liver T_RM_ cell numbers. Data were log transformed and statistically compared using two-way ANOVA and Tukey’s multiple comparisons test. **F.** Percentages of liver T_RM_ cells of different specificities amongst all T_RM_ cells in the liver. **G.** Fold change in the number of T_RM_ cells of the indicated specificities compared to those in 1xCS5M RAS vaccinated mice. Data were log transformed and statistically compared using one-way ANOVA and Tukey’s multiple comparisons test. Data in D-F were pooled from two independent experiments. **H.** Ratios of T_RM_ cell numbers in the liver vs numbers of circulating memory T cells (T_CIRCM_, calculated by adding T_CM_ and T_EM_ cells) in the spleen. Data were log-transformed and compared using unpaired Student’s t-tests.

To consolidate these findings, we next determined whether a similar bias occurred for T cells specific for sporozoite-associated antigens other than TRAP. We focused on a prototypical, abundant sporozoite antigen, the circumsporozoite protein (CSP). As C57BL/6 mice do not respond to *P. berghei* CSP, we utilised *P. berghei* CS5M parasites (termed CS5M henceforth) for RAS immunisations. In these parasites, CSP was mutated to encode the OVA_257-264_ (SIINFEKL) peptide, recognised by endogenous CD8^+^ T cells in C57BL/6 mice [43]. Mice were vaccinated with 1 or 3 doses of CS5M RAS and the numbers of memory CD8^+^ T cells specific for OVA, TRAP, RPL6 or RPA1 were measured 30 days later (Figure 2E-H, S3). OVA-specific, as well as TRAP specific T_RM_ cells, expanded to much higher numbers (approximately 46 and 51-fold increases respectively, Figure 2G) than RPL6- or RPA1-specific T_RM_ cells (3.5 and 1.6-fold increase respectively) in 3x vs 1x CS5M RAS vaccinated mice, with T_RM_ cells of other specificities increasing 4.3-fold (Figure 2E-G, S3). The combined frequencies of CS5M-CSP and TRAP-specific cells accounted for close to half of all liver T_RM_ cells (Figure 2F). Interestingly, in this case, OVA-specific T cells formed substantially higher numbers of memory cells (4-fold more on average) than those specific for TRAP (Figure 2E, F, S3), which did not expand to comparable numbers as when WT RAS were used for immunisations (Figure 2A) and hence no known, potent competing sporozoite-specific T cell response was elicited. This suggested that CS5M-CSP specific responses outcompeted those against TRAP, even though both specificities were preferentially expanded in comparison to RPL6 or RPA1. As observed previously (Figure 2D), for all specificities, repeated RAS vaccinations biased memory T cell formation towards liver T_RM_ cells over T_CIRCM_ cells (Figure 2H). Together, these results verified that multiple RAS vaccinations favoured the generation of resident memory CD8^+^ T cells specific for abundant sporozoite antigens over those responding to other antigens.

### Repeated RAS vaccination generates long lived liver T_RM_ cells

Having observed that repeated RAS vaccination altered the memory T cell phenotype by progressively favouring the development of T_RM_ cells, we next aimed to investigate whether boosting could modify other intrinsic properties of these cells. To this end, we assessed the impact of multiple RAS immunisations on the maintenance of liver T_RM_ cells of different specificities over time. Mice were injected with either one dose of 5x10^4^ RAS, or three doses of 10^4^ RAS administered weekly, and the numbers of TRAP-, RPL6- and RPA1-specific memory T cells were measured in the liver and spleen at several time points extending up to 100 days post-immunisation (Figure 3). Remarkably, T_RM_ cells of all specificities examined in mice vaccinated with 3xRAS displayed markedly increased half-lives (Figure 3A-C), though this trend was not significant for RPA-1 (p=0.1023). Specifically, the half-lives of liver T_RM_ cells in 1xRAS vaccinated mice ranged from 47 to 81 days, while those in 3xRAS vaccinated mice extended to nearly 230 days for TRAP-specific cells (Figure 3A), and exceeded a thousand days for the later antigens, RPL6 and RPA1 (Figure 3B, C). Interestingly, this increase in longevity was less pronounced and failed to reach significance for T_EM_ cells. For all specificities, spleen T_EM_ cells displayed much shorter half-lives than liver T_RM_ cells and exhibited moderate increased half-lives in 3xRAS mice, reaching 30-85 days, up from 20-50 days in 1xRAS mice (Figures 3D-F). In summary, these results showed that repeated RAS vaccination increased the life span of parasite-specific memory T cells, with liver T_RM_ cells displaying much larger increases.

**Figure 3.**
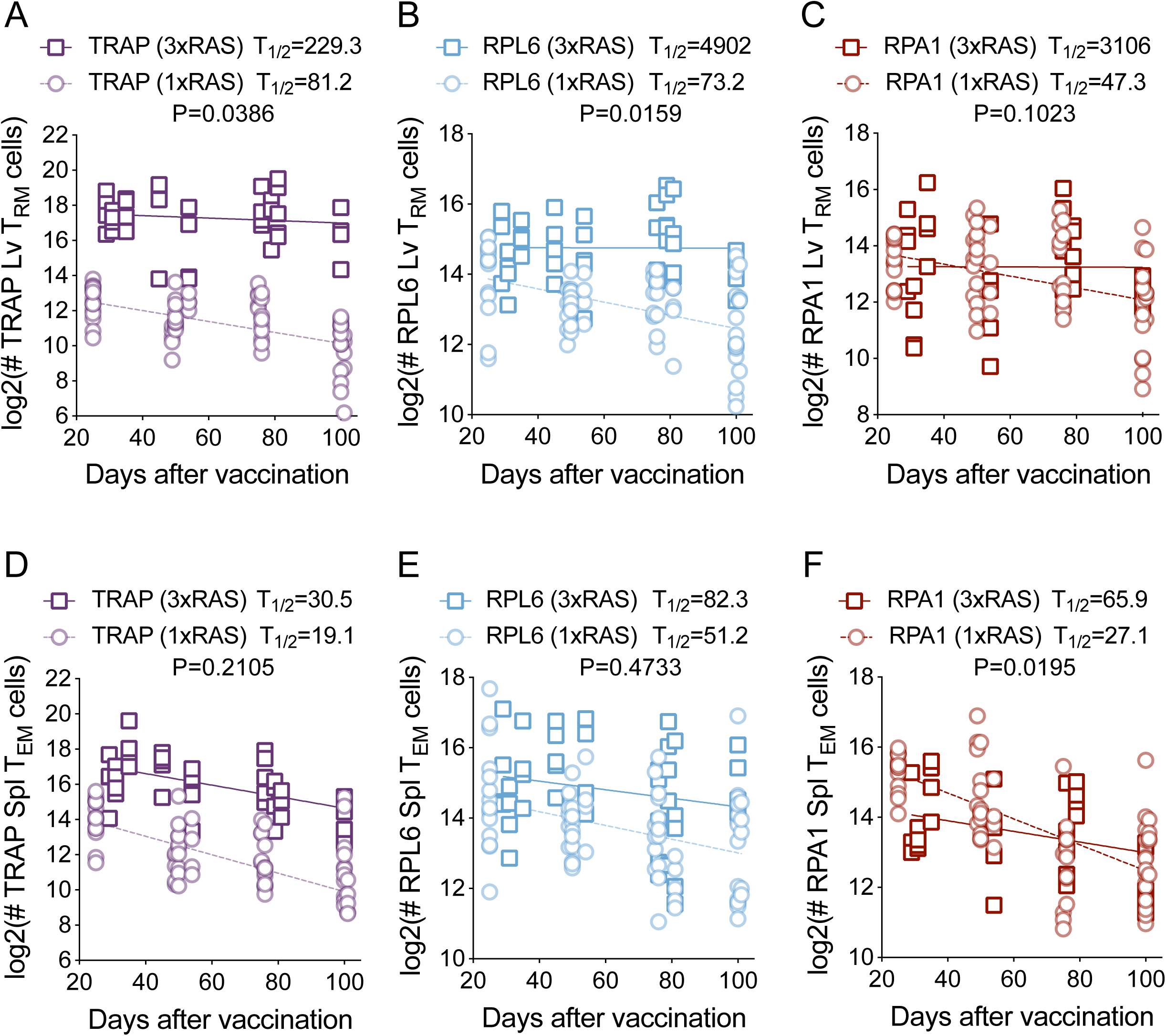
Repeated RAS vaccination generates long lived memory T cells. Numbers of CD8^+^ T_RM_ cells in the liver (**A-C**) or T_EM_ cells in the spleen (**D-F**) specific for TRAP (**A, D**), RPL6 (**B, E**) and RPA1 (**C, F**) in mice vaccinated with one dose of 50,000 RAS (dotted line) or three doses of 10,000 RAS (solid line), one week apart. Cell numbers were assessed up to day 101 after the last round of immunisation. Data were pooled from 5 independent experiments for 1xRAS and 8 independent experiments for 3xRAS. Data were log-2 transformed and linear regression analyses of the log-transformed data were performed. Slopes were compared using F-tests.

### TRAP-specific immunity dominates protection

To understand the implications of these findings, we sought to determine the contribution of TRAP-specific T_RM_ cells to protection against challenge with *P. berghei* sporozoites in RAS vaccinated mice. To do this, we suppressed the development of TRAP specific responses by sporozoite vaccination through injection of PbTRAP_130-138_ peptide in the absence of adjuvant, which leads to the removal of T cells specific for this peptide [40, 44]. Mice received 3 doses of TRAP peptide diluted in PBS prior to the first dose of RAS, and then received additional injections a day before administration of the second and third doses of the vaccine (Figure 4A). This resulted in efficient deletion of TRAP-specific cells (Figure S4A). In contrast, administration of an irrelevant peptide (SIINFEKL, i.e. OVA_257-264_) did not alter the generation of TRAP specific memory cells (Figure S4A). Vaccination with three doses of RAS induced strong protection against sporozoite challenge in untreated and SIINFEKL-treated control mice. However, TRAP-tolerised mice displayed markedly reduced levels of sterile protection, comparable to unvaccinated mice, and higher parasitemia 7 days after challenge (Figure 4B, C). Those TRAP-tolerised mice that became infected had significantly lower parasitemias than unvaccinated mice, suggesting that T_RM_ cells of other specificities contributed moderately to protection (Figure S4B). Enumeration of liver T_RM_ cells in challenged mice (Figure 4D) showed that, as expected, 3xRAS vaccination induced substantial numbers of TRAP-specific liver T_RM_ cells, except in the single mouse that remained unprotected (a second unprotected mouse could not be analysed as it developed cerebral malaria and had to be euthanised). TRAP tolerization efficiently impaired formation of TRAP-specific liver T_RM_ cells in unprotected mice, but two of the TRAP-tolerised mice that remained protected had high numbers of TRAP T_RM_ cells, indicating that, in these mice, tolerisation did not work efficiently (Figure 4D). Also, one out of the two non-protected mice in the OVA-tolerised control group tended to have low numbers of TRAP specific cells (similarly to the unprotected mouse in the RAS group), indicating that either RAS immunisation, or expansion of TRAP-specific cells, were suboptimal in this mouse (Figure 4D). Together, these results demonstrated a key role for TRAP-specific T_RM_ cells in protection against sporozoite challenge in C57BL/6 mice vaccinated multiple times with RAS.

**Figure 4.**
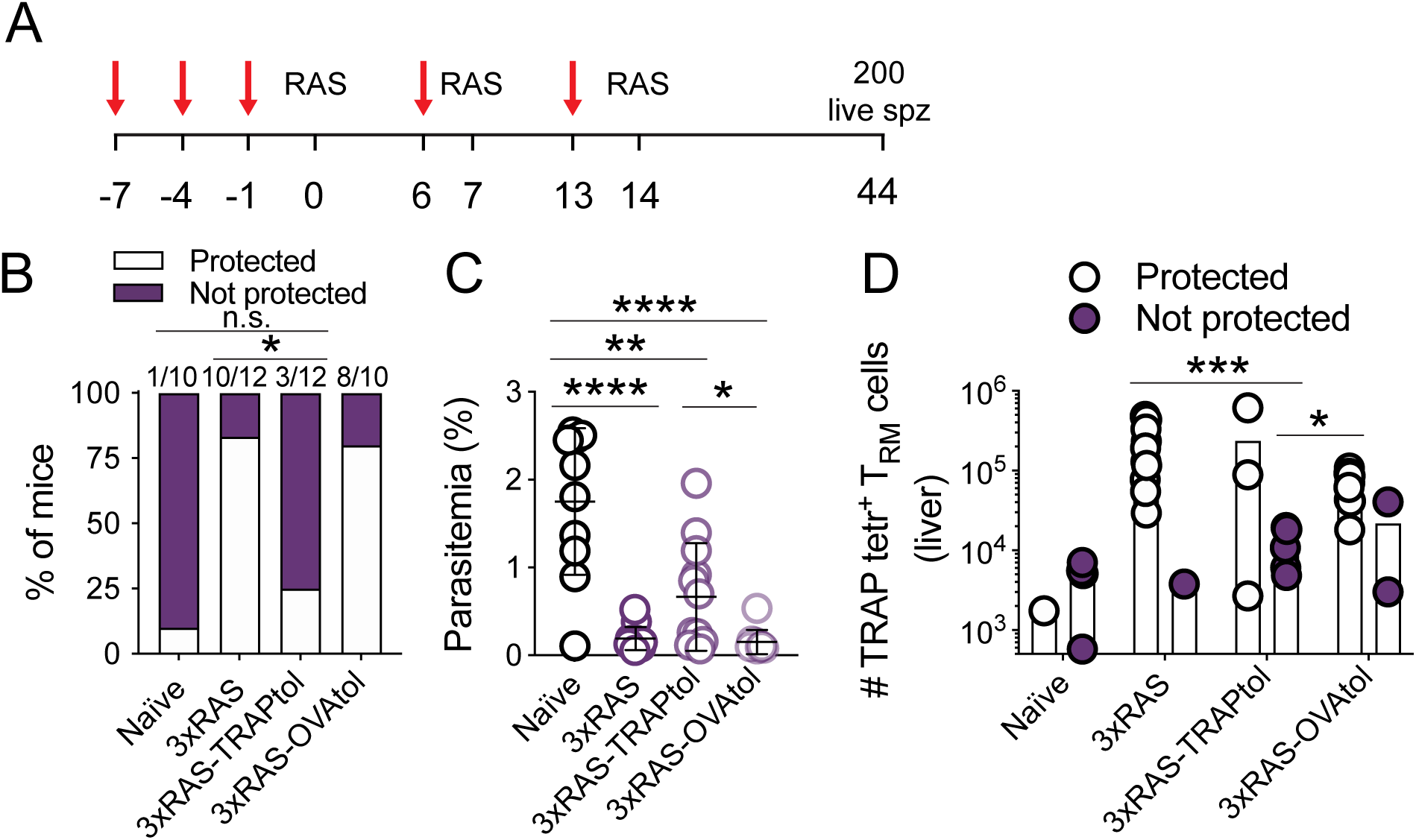
TRAP-specific liver T_RM_ cells substantially contribute to protection induced by 3xWT RAS in C57BL/6 mice. **A.** Experimental design. Red arrows denote intravenous injection of PbTRAP_130-138_ or OVA_257-264_ peptide dissolved in PBS. Numbers denote days after the first RAS vaccination. **B.** Sterile protection to live sporozoite challenge in 3xRAS vaccinated mice in which TRAP-specific cells (3xRAS-TRAPtol), or T cells specific for an irrelevant antigen (3xRAS-OVAtol) were removed (tol=tolerated). Data were compared using Fisher’s exact test. **C.** Parasitemia on day 7. Data were log-transformed and compared using one-way ANOVA and Tukey’s multiple comparisons test. **D.** Number of TRAP-specific liver CD8^+^ T_RM_ cells in protected vs non-protected mice. Data were log-transformed and compared using two-way ANOVA and uncorrected Fisher’s LSD test. Data in this figure were pooled from two independent experiments.

### RAS preferentially boosts previously activated T cells

A potential mechanism to explain the T cell memory bias towards sporozoite antigen specificities generated by repeated RAS vaccination was the rapid elimination of incoming irradiated sporozoites by CD8^+^ T cells induced by prior doses of RAS. This elimination would particularly curtail the expression of late antigens by booster RAS, thereby limiting their immunogenicity compared to the readily available antigens present in the sporozoite [45]. To test whether prior RAS vaccination limited the development of parasites subsequently administered in booster doses, mice were vaccinated with one or two consecutive doses of RAS up to 8 days apart. Then, coinciding with the timing of the final dose in our established RAS vaccination schedule, mice were instead challenged with an equivalent dose (10,000 parasites) of non-irradiated, fully infectious sporozoites. We then assessed protection to evaluate whether the pre-existing immune response interfered with the development of these live sporozoites (Figure 5A). Mice that received a single dose of RAS prior to infection displayed marked decreases in parasitemia on day 7 compared to unvaccinated controls, and some sterilising protection (Figure 5B, C). Moreover, two doses of RAS provided complete sterilising protection (Figure 5B, C). In agreement with previous work [45], these results indicated that the effector response elicited by prior doses of RAS has the capacity to efficiently eliminate incoming sporozoite infections, potentially impairing the generation of T cell responses against liver stage antigens upon administration of subsequent doses of RAS.

**Figure 5.**
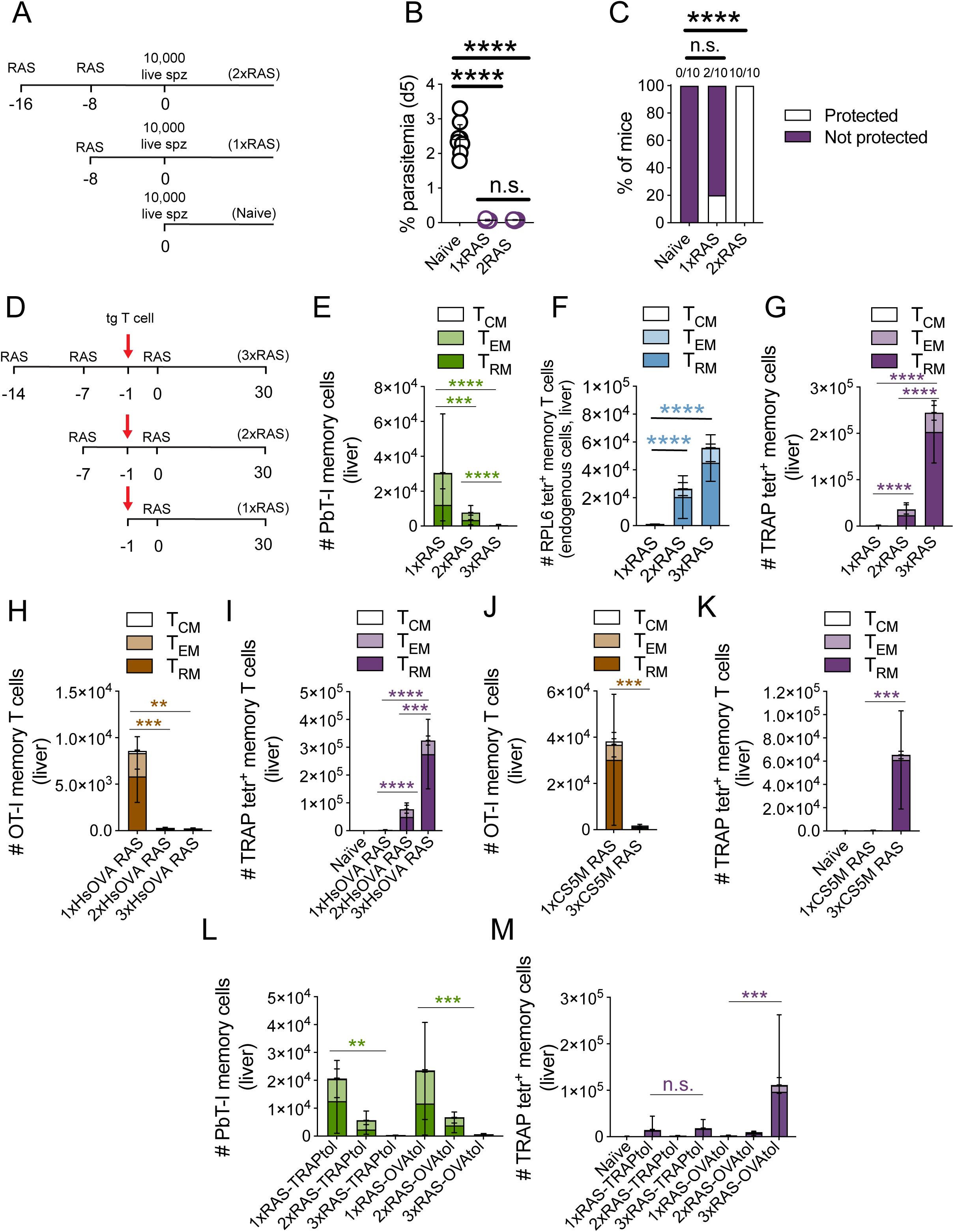
Mechanisms contributing to T cell specificity bias towards sporozoite antigens. **A-C.** Protective capacity of effector responses elicited by RAS against incoming sporozoites. **A.** Representative experimental design. **B-C.** Mice received one or two doses of 5-10x10^3^ RAS 4-8 days apart, and 8-9 days later were injected with 1x10^4^ live sporozoites. Emergence of parasitemia was measured to evaluate protection. Data were pooled from two independent experiments. **B.** Parasitemia on day 5 after liver sporozoite challenge. Data were log-transformed and compared using one-way ANOVA and Tukey’s multiple comparisons test. **C.** Sterile protection. Data were compared using Fisher’s exact test. Numbers above columns denote numbers of protected mice / total numbers of mice per group. **D-K.** Inhibition of naïve CD8^+^ T cell responses by prior RAS vaccination. **D.** Representative experimental design. Red arrows denote the time point in which mice were intravenously injected with 5x10^4^ naïve transgenic T cells (PbT-I cells in D-F; OT-I cells in F-L). **E-G.** Inhibition of naïve RPL6-specific (PbT-I) T cell responses by prior RAS vaccination. Distribution of memory PbT-I (**E**) and endogenous, RPL6-(**F**) and TRAP-specific (**G**) T cells in the liver 30-35 days after the last RAS injection. Mice received 1-3 doses of 5-10x10^3^ RAS. Data were pooled from two independent experiments. Statistical analyses, denoted by dark green and purple asterisks, denote comparisons of T_RM_ numbers. **H-I.** Inhibition of naïve HsOVA-specific (OT-I) T cell responses by prior HsOVA RAS vaccination. Distribution of memory OT-I (**H**) and endogenous, TRAP-specific (**I**) T cells in the liver 30 days after the last HsOVA RAS injection. Mice received 1-3 doses of 5.9-10x10^3^ HsOVA RAS. Data were pooled from two independent experiments. Statistical analyses, denoted by brown and purple asterisks, denote comparisons of T_RM_ numbers. **J-K.** Inhibition of naïve CSP-specific (OT-I) T cell responses by prior CS5M RAS vaccination. Distribution of memory OT-I (**J**) and endogenous, TRAP-specific (**K**) T cells in the liver 30 days after the last CS5M RAS injection. Mice received 1 or 3 doses of 5.9-10x10^3^ CS5M RAS. Data were pooled from two independent experiments. Statistical analyses, denoted by brown and purple asterisks, denote comparisons of T_RM_ numbers. **L-M**. Naïve CD8^+^ T cell responses to RPL6 are inhibited in the absence of TRAP-specific T cells. Mice were vaccinated with 1-3 doses of 5-10x10^3^ WT RAS and received intravenous injections of either PbTRAP_130-138_ (TRAPtol) or OVA_257-264_ peptide (OVAtol) dissolved in PBS as explained in Figure 4A. PbT-Is were transferred 1 day before the last RAS vaccination and, on day 36 after the last RAS vaccination, mice were euthanised and numbers of memory PbT-I or TRAP-specific memory CD8^+^ T cells were examined in the liver. **L.** Number of memory PbT-I cells in the liver. **M.** Numbers of TRAP-specific memory T cells in the liver. Data were pooled from two independent experiments, log-transformed and analysed using two-way ANOVA and Tukey’s multiple comparisons test.

Given the strong bias towards the development of sporozoite-specific responses observed in figure 2, we reasoned that not all T cell responses elicited by RAS would be equally impaired by this inhibitory mechanism. Particularly, we anticipated that responses targeting abundant antigens in the incoming sporozoites would be less affected by quick parasite killing that those specific for later or less abundant antigens. To test this hypothesis, we sought to define the capacity of RAS to activate naïve T cells of different specificities for parasite antigens, in mice that had been previously vaccinated with RAS, and hence had an ongoing T cell response capable of killing further incoming parasites. In a first series of experiments, we adoptively transferred naive PbT-I cells, i.e. RPL6-specific TCR transgenic T cells [31, 37], into mice that had received no prior RAS vaccination, or one or two doses of RAS one week apart. Mice then received a final dose of RAS one day after PbT-I T cell transfer, and the numbers of memory cells generated by these transgenic T cells were examined 30 days after the last RAS injection (Figure 5D). As expected, when PbT-I cells were transferred one day before a single RAS vaccination, substantial numbers of memory PbT-I cells formed in the spleen and the liver, including T_RM_ cells (Figure 5E, S5A). However, when naïve PbT-I cells were transferred one day before a second RAS vaccination, fewer PbT-I liver T_RM_ cells formed. Furthermore, when transferred prior to the third RAS immunisation, virtually no memory PbT-I cells were detected (Figure 5E, S5A). Importantly, although PbT-I responses were strongly inhibited, endogenous memory CD8^+^ T cells of the same specificity (i.e. RPL6-specific) increased in numbers upon successive RAS vaccinations (Figure 5F, S5A), and the same occurred for RPA1-specific cells (Figure S5A). As observed before, large numbers of TRAP T_RM_ cells formed in these mice (Figure 5G, S5A). Note that, in agreement with previous reports [37], PbT-I cells did not detectably respond to persisting antigen when adoptively transferred into mice vaccinated with RAS 6 days earlier (Figure S5B), and therefore the observed responses were induced by RAS administered after adoptive T cell transfer. At an early time point (7 days) after the last injection of RAS, PbT-I cells were virtually undetectable in the blood of mice receiving several doses of RAS prior to PbT-I transfer, while numbers of endogenous TRAP-or RPL6-specific T cells increased in the same mice upon administration of every additional dose of RAS (Figure S5C). This indicated that diminished early expansion of PbT-I cells was responsible for the low numbers of memory PbT-I cells subsequently detected. Moreover, this occurred while endogenous CD8^+^ T cells of the same specificity, activated by prior RAS vaccination, continued expanding.

To extend these findings and better understand the mechanisms contributing to sporozoite antigen immunodominance, we next sought to determine whether this inhibitory effect also applied to other T cell specificities. To do this, we performed similar experiments as those explained for PbT-I cells, but instead using naïve T cells of a different specificity, i.e. OVA-specific OT-I cells [46]. To provide a target antigen for these cells, we utilised *P. berghei* HsOVA (termed HsOVA henceforth) parasites for vaccination. In these parasites, a C-terminal fragment of OVA (aa 150-386) was fused to the N terminus of a truncated version of the parasite protein Hsp70 (retaining aa 201-398) [47]. This construct is expressed under the control of the Hsp70 promoter, which induces high levels of transcription in sporozoites and during the liver stage (Figure S5D) [30]. However, as mRNA is not necessarily indicative of protein expression [48] or immunogenicity, we first sought to determine the T cell immunogenicity of this antigen in sporozoites. Mice received OT-I cells and were intravenously injected with heat killed sporozoites (HKS), thereby limiting the repertoire of presentable antigens to T cells to those already existing in the sporozoite. Expansion of OT-I cells was assessed in the spleen after 4 days. PbT-I cells were utilised as positive controls, as they can respond to HKS [49]. We found that, indeed, OT-I cells moderately responded to HsOVA HKS (Figure S5E), confirming the immunogenicity of this antigen in sporozoites. We then proceeded to assess whether repeated RAS vaccination resulted in the inhibition of naïve T cell responses specific for Hsp70-OVA. As with the previous experiments with PbT-I cells, mice were vaccinated 1-3 times with HsOVA RAS, and naïve OT-I cells were transferred one day prior to the last HsOVA RAS vaccination. Although OT-I cells generated strong memory in 1xHsOVA RAS vaccinated mice, this response was completely inhibited in mice previously vaccinated once or twice with HsOVA RAS (2xHsOVA RAS and 3xHsOVA RAS respectively) (Figure 5H, S5F), appearing even more strongly inhibited than the response by PbT-I cells (Figure 5E). As expected, endogenous TRAP specific cells formed substantially increasing numbers of T_RM_ cells upon every additional RAS injection (Figure 5I, S5F), with RPL6-specific T_RM_ cells, as well as T_RM_ cells of undefined specificity, displaying more moderate increases (Figure S5F).

### Preferential boosting of pre-activated T cells extends to abundant sporozoite antigens

We hypothesised that suppression of naïve responses in this system would be less pronounced if the adoptively transferred naïve T cells were specific for a readily available, abundant sporozoite antigen, such as CSP (Figure S6A). To test this, we examined the expansion of naive CD8^+^ T cell responses targeting CSP in mice previously vaccinated with RAS. Here, we again used CS5M RAS, where the OT-I epitope is embedded within CSP and OT-I cells can be used as a readout for responses to *P. berghei* CSP. To demonstrate the immunogenicity of CS5M-CSP expressed on these sporozoites, we adoptively transferred naïve OT-I cells into recipient mice, which were injected with heat killed CS5M sporozoites a day later. Expansion of OT-I cells was then assessed in the spleen after 6 days. CS5M HKS induced substantial OT-I proliferation, indicating that sporozoite-derived surface CS5M-CSP could be captured and presented to responding T cells by host antigen presenting cells (Figure S6B). In a new set of experiments, mice then either received a single injection of CS5M RAS, or three injections, with OT-I cells being transferred one day before the last RAS vaccination. Surprisingly, *de novo* OT-I responses in 3xCS5M RAS vaccinated mice were again strongly suppressed (Figure 5J), even when endogenous SIINFEKL-specific responses strongly expanded (Figure S6C). TRAP specific, as well as RPL6 and T cells of undefined specificities, formed large numbers of memory cells in 3xRAS vaccinated mice (Figure 5K, S6D). Moreover, mice that, in a separate experiment, received two doses of WT RAS, followed by OT-I transfer before a last dose of CS5M RAS, in which therefore no OVA specific response was generated by prior to CS5M RAS vaccination, also displayed strongly diminished OT-I T_RM_ cell numbers (Figure S7). These results confirmed that naive T cell responses are strongly inhibited by ongoing responses elicited by prior RAS vaccination. This is the case, even for responses specific for an abundant sporozoite antigen that can be cross-presented without requiring liver invasion (Figure S6B), and can even occur when the challenge antigen has not been encountered during previous vaccinations. Furthermore, this inhibition of naïve T cell responses occurs while previously activated T cells of the same specificity continue to respond to each RAS vaccination.

### Inhibition of naïve T cell activation by RAS boosting does not require TRAP-specific T cells

Having established that TRAP-specific T cells dominate the protective response elicited by multiple WT *P. berghei* RAS vaccinations in C57BL/6 mice, we sought to determine whether these cells were also responsible for the observed inhibition of naïve T cell responses. To do this, we transferred naïve PbT-I cells into mice vaccinated with 1x, 2x or 3xWT RAS, one day before the last RAS vaccination, as done before, but in this case, vaccinated mice were tolerised for TRAP or OVA prior to RAS vaccination as done before (Figure 4A). Memory formation by PbT-I cells was measured 30 days after the last RAS vaccination (Figure 5L, M, SF8). Naïve PbT-I cells were impeded from forming memory (Figure 5L, S8A), despite TRAP tolerisation having efficiently removed the TRAP-specific response in most mice (Figure 5M, S8B). As observed in previous experiments (Figure 4D), TRAP tolerization did not work efficiently in all mice, yet a clear inhibition of PbT-I responses could be observed in those mice in which TRAP tolerization was efficiently achieved (Figure S8C, D). T_RM_ cells of undefined specificities (other than RPL6 or TRAP) progressively increased in numbers after each dose of RAS (Figure S8E, F), independently of the presence or absence of TRAP specific cells. These results show that, although TRAP specific cells contribute decisively to protection, inhibition of naïve T cell responses in 3xRAS mice was not exclusively mediated by effector T cells of this specificity. Taken together, these data suggested that immunity generated after the initial RAS immunisation hindered the boosting of effector CD8^+^ T cell responses to late epitopes, and dramatically inhibited the priming of new T cell responses to any antigen, either sporozoite or liver stage derived. Overall, this resulted in a strong bias of the T cell response towards sporozoite-derived immunogenic antigens associated with the primary exposure to RAS.

### Liver T_RM_ cells specific for abundant sporozoite antigens can confer potent protection

To better understand the implications of the skewed T_RM_ specificities towards sporozoite proteins, we next sought to explore the protective capacity of liver T_RM_ cells specific for the major sporozoite antigen CSP, and benchmarked this protection with that provided by T_RM_ cells specific for Hsp70. This was achieved by evaluating the ability of OT-I T_RM_ cells to protect against challenges with either CS5M or HsOVA parasites. This experimental design allowed us to directly compare the efficacy of T_RM_ cells of the same specificity (SIINFEKL) against two modified antigens containing this epitome, namely CS5M-CSP and Hsp70-OVA. As reported above, Hsp70 is present in the sporozoite and, although it is moderately immunogenic at this stage (Figure S5D), it is prominently transcribed during liver stage (Figure S5C) and can be targeted for protection [50]. In turn, CSP combines abundant expression and immunogenicity in sporozoites (Figure S6A) with prolonged expression during liver stage [51]. To ensure protective responses were only specific for the relevant OVA peptide, SIINFEKL, expressed by HsOVA and CS5M parasites, and not for other parasite antigens, we generated OVA-specific responses using our previously developed prime-and-trap (P&T) approach [22]. Mice either received OT-I cells or not (and hence relied on endogenous SIINFEKL-specific CD8^+^ memory T cells for protection), and were challenged with CS5M or HsOVA parasites. Large numbers of OT-I T_RM_ (Figure 6A-C), or SIINFEKL-specific endogenous cells (Figure 6D-F), were generated in mice receiving the complete P&T vaccine. When these mice were challenged with CS5M or HsOVA parasites, high levels of sterile protection were achieved (Figure 6G, H). As T_RM_ cell numbers generated with this vaccination strategy provided exceedingly efficient protection against both parasites, we decided to vaccinate mice suboptimally, to better gauge the numbers of T_RM_ cells required for protection against these parasites. Thus, we injected mice with a low dose of Clec9A-OVA (0.5 μg) plus 5 nmol CpG B-P adjuvant. This induced lower numbers of liver OT-I T_RM_ cells, averaging 5x10^4^ cells (Figure 6I). These mice displayed substantial sterile protection against CS5M CSP, which was superior to that of Hsp70-OVA (Figure 6J). This result showed that T_RM_ cells specific for sporozoite antigens, particularly those abundantly expressed, can provide highly efficient protection against infection. Overall, this supported the view that bias towards sporozoite-specific T_RM_ cells induced by repeated RAS vaccination provided highly efficient protection against infection.

**Figure 6.**
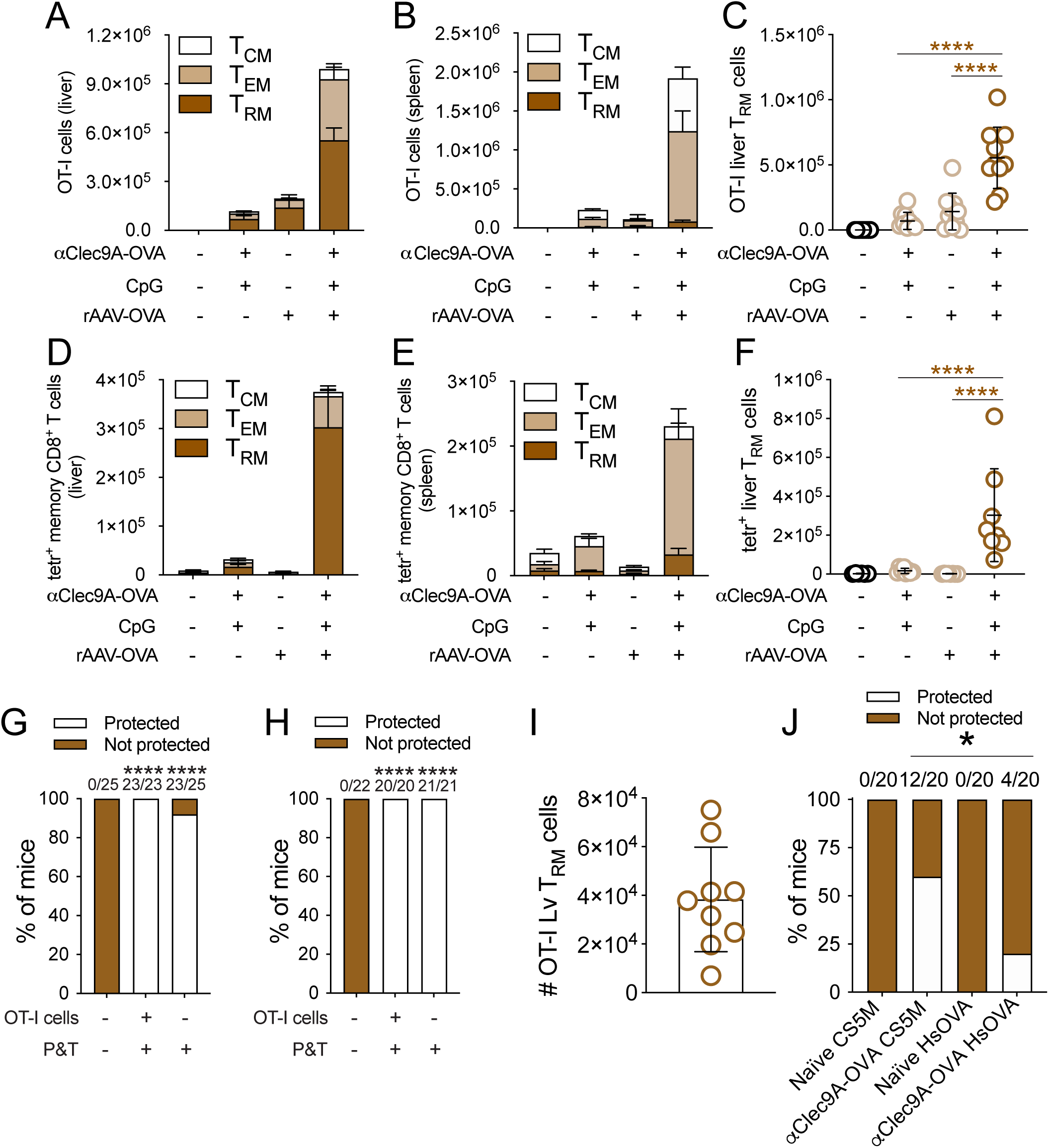
Memory T cell generation of prime-and-trap targeting OVA, and protection against challenge with parasites expressing SIINFEKL under the Hsp70 promoter or within CSP. **A-H**. Mice received 50,000 naïve OT-I/uGFP cells (**A-C, G, H**) or not (**D-F, G, H**) and were vaccinated with either 2 μg Clec9A-OVA plus 5 nmol CpG B-P, 10^9^ rAAV-OVA, or all components combined. OT-I (**A-C**) or SIINFEKL-specific memory cells (**D-F**) were enumerated in the liver and the spleen 35 days later. Data were pooled from two independent experiments, log transformed and compared using one-way ANOVA and Tukey’s multiple comparisons test. Mice vaccinated with full P&T-OVA were challenged with 200 HsOVA (**G**) or CS5M (**H**) sporozoites on day 35 after vaccination, and rates of sterile protection were determined. Data were pooled from two independent experiments and compared using Fisher’s exact test. Asterisks over columns denote comparisons with the unvaccinated control group. Numbers over columns denote numbers of protected mice vs total numbers of mice. **I-J.** Generation of OVA-specific memory T cells in suboptimally vaccinated mice. Mice received 50,000 naïve OT-I/uGFP cells and were vaccinated with a low dose of 0.5 μg Clec9A-OVA plus 5 nmol CpG B-P. Numbers of OT-I T_RM_ cells were measured in the liver 30 days later (**I**). Separate cohorts of mice were challenged with 200 HsOVA or CS5M sporozoites (**J**), and rates of sterile protection were determined. Data were pooled from two independent experiments and compared using Fisher’s exact test.

Together, this study shows that repeated RAS vaccination enhances the development of long-lived liver-resident memory CD8^+^ T cells of biased specificity for abundant sporozoite antigens, some of them highly protective, but fails to fully exploit the protective potential of T cells specific for less abundant or later antigens.

## Discussion

RAS can induce efficient protection against malaria across a diverse range of species, including humans [9, 13], non-human primates, and mice. The rodent model provides the opportunity to expand our understanding of this vaccine, as well as to investigate fundamental immunology processes, to inform the design of more effective malaria vaccines.

Once inside hepatocytes, malaria parasites become targets of CD8^+^ T cell immunity. The pivotal role of memory CD8^+^ T cells in protecting mice [10, 14, 15] and NHP [16] during the liver stage of malaria has been well established. However, not all types of memory CD8^+^ T cells are equally efficient at providing protection. We here show that liver T_RM_ cells, which afford highly focused and efficient responses to infection [22], are critical mediators of the immunity to *P. berghei* conferred by this vaccination strategy in C57BL/6 mice. Existing data also supports a prominent protective role for liver CD8^+^ T_RM_ cells in primates. CD8^+^ T cells had been observed to accumulate in the liver in NHP vaccinated with RAS [19], prior to the identification of liver T_RM_ cells. In humans, protection was maintained in RAS-vaccinated individuals a year after vaccination, a time when numbers of circulating T cells and antibodies had declined to background levels [13]. Further studies showed that numbers of parasite-specific circulating CD8^+^ T cell numbers generated by RAS are low, and do not discriminate between protected and non-protected individuals [19, 25, 52]. The results presented here mirror these observations, as repeated RAS vaccination in mice does not prominently increase numbers of circulating memory cells but comparatively enhances liver T_RM_ numbers and protection. Additionally, protection increases with successive RAS injections, mirroring observations in humans [9]. This mouse model hence presents strong analogies with humans and is a valuable tool to study immunity to pre-erythrocytic malaria in the liver.

T_RM_ cells are more abundant than circulating memory T cells [53], and have been identified and comprehensively characterised in humans as analogous to those in mice [54]. These cells express high levels of cytokines and cytotoxicity mediators [22, 55, 56] and exert rapid effector function [57, 58] for potent, protective responses against infection [22, 59-62]. The significant protective capacity of liver T_RM_ cells against malaria has been previously demonstrated [22, 63, 64], although circulating, effector memory CD8^+^ T cells also infiltrate the liver and were shown to contribute to protection against *P. berghei* infection in CB6F1 mice [21]. In C57BL/6 mice, which are highly susceptible to infection by *P. berghei* [23], liver T_RM_ cells appear to be critical for protection, with T_CIRCM_ cells failing to provide substantial protection even at considerable numbers [22]. In line with these findings, we here show that depletion of T_RM_ cells rendered RAS vaccinated mice devoid of sterile protection, and the parasitemia developed by depleted mice, although slightly lower, was statistically comparable to that of unvaccinated mice. Additionally, the increase in protection observed upon administration of successive doses of RAS was associated with a comparably more pronounced increase in T_RM_ cells than T_CIRCM_ cells, as exemplified by the improved ratio of the former cells vs the latter. As T cell responses derived from naïve T cells are strongly inhibited in repeatedly vaccinated mice (discussed below), the observed increase in the T_RM_/T_CIRCM_ ratio may be due to local proliferation of liver T_RM_ cells upon antigen encounter [65, 66], or to conversion of T_CIRCM_ cells into T_RM_ cells [67, 68]. In line with our findings, repeated immunisation with *Listeria monocytogenes* has been previously found to progressively increase the accumulation of specific memory T cells in the liver [69]. Together, our findings showcase the capacity of RAS to generate T_RM_ immunity and underscore a major role of these cells in the induced protection against malaria.

Despite their intrinsic ability for efficient pathogen control, T_RM_ cells are constrained by their antigen specificity in their capacity to provide protection against infection. Thus, defining the specificities that drive immunity in RAS vaccinated mice is critical to understand the mechanism of protection of this vaccine, and can be invaluable for antigen selection for subunit vaccine development. Upon invasion of the liver as sporozoites, parasites proliferate massively within hepatocytes and progressively develop towards the merozoite form, able to invade erythrocytes. This process entails pronounced variations in protein expression [41, 51, 70], and therefore in available immune targets. Vaccination with genetically modified parasites engineered to interrupt their intrahepatic development at a late stage after hepatocyte invasion induces more protective T cell responses than early arresting parasites, which express a more limited antigen breadth [33]. Similarly, as RAS parasites die quickly upon hepatocyte invasion [8], the repertoire of antigens they present to the immune system is limited [33]. In addition to inducing responses of comparably lower protective quality, the logical consequence of this phenomenon, as we have observed, is that the expansion of T cells specific for early antigens, such as those present in the sporozoite, is favoured, introducing a bias for these specificities in the T_RM_ cell repertoire generated. Our results parallel prior observations in RAS-vaccinated BALB/c mice [45], where responses specific for a sporozoite antigen (CSP), but not those recognising a liver stage antigen, were enhanced through repeated *P. yoelii* short-interval RAS immunisations. Importantly, *P. yoelii* CSP features an immunodominant K^d^-restricted epitope in BALB/c mice [71], which hence becomes a clear target of T cell immunity, whereas C57BL/6 mice do not mount CD8^+^ T cell responses to *P. berghei* CSP [36]. As we have found here, the response elicited by RAS in this mouse strain is instead dominated by T cells specific for TRAP, another major sporozoite antigen [41, 72]. Although expected on the basis of the evidence presented above, this is particularly striking because TRAP is poorly recognised by the CD8^+^ T cell compartment of C57BL/6 mice, with only about 1 cell per million naïve CD8^+^ T cells specific for its single known MHC-I-restricted epitope [31, 73]. Indeed, when *P. berghei* CSP is modified to contain SIINFEKL, a much more immunogenic antigen in C57BL/6 mice [35], and therefore both CSP- and TRAP-derived CD8^+^ T cell antigens are concurrently present in the same, early sporozoite stage at approximately similar levels [41], then CSP-specific cells clearly outcompete TRAP specific cells (Figure 2D, E), evidencing the comparative weakness of the latter antigen. Nevertheless, TRAP-specific cells expand to much greater numbers than RPL6-specific cells in mice vaccinated with three doses of WT RAS, even though the frequency of the latter cells in the naïve repertoire is about 100-fold higher [31]. These results underline the surprising strength of the bias towards T cells specific for early antigens induced by RAS.

A straightforward explanation for this phenomenon is that proliferation of parasite-specific T cells is strongly dictated by early antigen availability, whereby T cells specific for abundant sporozoite antigens are preferentially expanded. Premature death, or quick elimination, of RAS parasites after hepatocyte invasion hence disfavours the expression and immunogenicity of later epitopes. Indeed, Murphy *et al.* monitored parasite mRNA expression in mice repeatedly vaccinated with RAS and found reduced mRNA expression of non-sporozoite antigens [45]. In line with these findings, we found that existing immunity generated during initial RAS vaccination strongly reduced the infectivity of subsequent sporozoite infections, potentially limiting the immunogenicity of later doses of RAS and likely reinforcing the T cell bias towards readily available early antigens. Possible sources of sporozoite antigens for CD8^+^ T cell immunity at early stages of infection are the parasite proteins that leak to the cytoplasm of the invaded hepatocyte and are presented via MHC-I molecules, as well as dying parasites that do not reach the liver and are captured by antigen presenting cells in lymphoid organs [74]. Although parasites can partially develop and express later antigens in these organs [32], we injected RAS intravenously, and therefore parasite development outside the liver is unlikely to have occurred [75]. Additionally, several sporozoite proteins, including CSP and TRAP, are cleaved and released as the parasites traverse hepatocytes [76, 77], generating a “gliding trail” of potential antigens that could potentially be captured by antigen presenting cells, or presented via MHC-I by the traversed hepatocytes. Moreover, those sporozoites that either die while traversing hepatocytes, or inside the final hepatocyte in which they settle for further development [78], may also become sources of additional antigen. We and others observed that HKS immunisation is markedly less immunogenic than RAS [19, 49, 74] and, although heat-induced damage to sporozoite proteins may explain these results, it is also possible that the simple cross-presentation of antigen that is readily available in sporozoites is not sufficient to induce potent immunogenicity. Liver invasion, or the presence of metabolically active parasites, may be required for maximal expansion of sporozoite-specific CD8^+^ T cell responses.

As mRNA levels can be poor indicators of immunogenicity, we attempted an alternative method to more accurately measure the immunogenicity of T cell antigens of interest with contrasting expression patterns. We did this by evaluating the memory formation capacity of adoptively transferred naïve TCR transgenic T cells in mice previously vaccinated with RAS. This led to our identification of another factor strongly contributing to sporozoite-specific T_RM_ cell bias, namely the strong inhibition of naïve T cell responses by prior RAS immunisation. This inhibition occurred progressively and was broad. It affected responses to all antigens examined, including those targeting abundant sporozoite antigens available for cross-presentation such as CSP, even when these antigens had not been encountered previously. Moreover, removing TRAP-specific responses, although significantly impacting protection, did not prevent inhibition of naïve T cell expansion. Notably, while naïve PbT-I cell activation was strongly inhibited, endogenous cells of the same specificity (RPL6), activated after the first RAS injection, continued to expand, albeit moderately, upon subsequent RAS injections (Figure S5A). Similarly, naïve OT-I cell expansion was hindered in CS5M RAS vaccinated mice (figure 3C) while the endogenous SIINFEKL-specific response continued to grow (Figure S6C). These findings suggested that this inhibition was not necessarily exerted by sporozoite-specific T cells over those specific for late antigens, but potentially by effector cells activated upon previous RAS injections, which outcompeted naïve T cells. Naïve CD8^+^ T cells have the capacity to proliferate massively over a few days following activation [38]. However, as pronounced changes in the naïve T cell, including a significant increase in size and a switch to glycolytic metabolism, must occur to enable such fast expansion, a period of about 24 hours is needed before proliferation commences [79]. This places naïve T cells at a disadvantage compared to already activated and expanding effector T cells. Additionally, activated T cells require factors such as IL-2 that enable further proliferation and subsequent formation of memory [80], and the more numerous effector T cells likely outcompete naïve T cells for these factors. Our results mirror those obtained by Hafalla *et al*, who observed that naïve CD8^+^ T cells specific for *P. yoelii* CSP expanded poorly when injected into mice immunised with a single dose of RAS 1-4 days earlier [81]. Subsequent work showed that competition for antigen presenting cells was another mechanism whereby effector T cells outcompeted naïve T cells [82]. Interestingly, this work observed that inhibition of naïve T cell responses by effector T cells only occurred when both responses were of the same specificity. However, we here present evidence that this suppression is broad and not restricted to specific antigens. In contrast to the scenario presented here, Hafalla *et al* utilised two different infection systems to create a competition between effector and naïve T cells (namely RAS and influenza infection). These systems may differ in parameters such as antigen availability or strength or quality of inflammatory signals, which could influence whether a *de novo* response can form. In summary, the effector T cell response generated upon primary RAS exposure constrains the breadth of the parasite-specific T_RM_ compartment generated by this vaccine, contributing to the establishment of a strong immunodominance biased towards early antigens that influences the quality of the protective liver T_RM_ response elicited by multiple RAS vaccinations.

Our work establishes sporozoite proteins as dominant antigens for naturally occurring T cell immunity to malaria in the liver. Notably, removing the TRAP response in 3xRAS vaccinated mice results in loss of protection. This finding was unexpected, given that 2x10^5^ TRAP T_RM_ cells generated by prime-and-trap vaccination, similar in number to those in 3xRAS mice, only conferred around 15% sterile protection against challenge with 200 *P. berghei* sporozoites in C57BL/6 mice [31]. This discrepancy could be attributed to the fact that RAS vaccination generates T_RM_ cells of multiple specificities [33], which can collectively contribute to protection to variable extents (Figure S4B), hence lowering the numeric requirements for TRAP-specific cells for sterilising protection. When a dominant specificity such as TRAP is removed, then the total number of protective T_RM_ cells is reduced below the threshold required for sterile protection, rendering mice susceptible to infection. Our results contrast with observations that substitution of *P. berghei* CSP for its ortholog in *P. falciparum*, which removes an immunodominant K^d^-restricted epitope in the rodent parasite [83], does not abrogate protection of BALB/c mice against challenge with WT sporozoites [84]. BALB/c mice are inherently more resistant that C57BL/6 mice to *P. berghei* sporozoite infection [18, 23], and can be efficiently protected by a comparably lower number of parasite-specific CD8^+^ T cells [18]. Hence, a potential explanation is that the pool of minor T_RM_ specificities suffice to exert protection in these mice.

Regarding further antigens, we found SIINFEKL-specific responses targeting Hsp70-OVA to be highly protective. This aligns with previous reports comparing vaccination targeting OVA expressed under the liver stage UIS4 promoter, and CSP, which found both antigens to be protective [85], yet we could determine by suboptimal vaccination that CS5M CSP responses induced more efficient protection than those against Hsp70-OVA. Comparatively strong protection by CD8^+^ T cells targeting sporozoite antigens has also been found by other groups [86]. However, the intrinsic protective capacity of an antigen depends on several factors, and not exclusively on its expression pattern. Thus, other immunogenic antigens with diverse expression patterns, including in the sporozoite, such as GAP50 or S20, or the liver stage, such as RPL3, were found not to be protective [40, 45, 86], a critical element for some of these antigens being antigen availability for MHC-I presentation in the hepatocyte [86].

Another important aspect of repeated RAS vaccination identified in this work is the extension of the lifespan of liver T_RM_ cells. Our results align with those obtained by van Braeckel-Budimir *et al.* on lung T_RM_ cells, whose lifespan is also significantly enhanced upon repeated influenza infection [68]. In that model, T_EM_ cells convert into T_RM_ cells after infection [67], themselves progressively decreasing in numbers [68], and consequently maintaining the T_RM_ compartment. Conversion of T_EM_ cells into T_RM_ cells has also been observed in the skin of Herpes Simplex virus infected mice [65], and a decrease in the lifespan of T_EM_ cells was also detected upon repeated stimulation with attenuated *Listeria monocytogenes* [87]. Our observation that lifespan extension is induced in T_RM_ cells, but not in circulating memory cells, is compatible with the occurrence of a similar phenomenon in our system. As naïve T cell responses are strongly inhibited by repeated RAS vaccination, this potential source of new T_RM_ cells is likely minor. In addition to recruitment from circulating memory cells, expansion of the T_RM_ compartment can also be due to *in situ* proliferation in the tissue in response to repeated antigen encounter [65]. Finally, repeated antigen stimulation induces intrinsic changes in memory T cells [69], including moderate changes in molecules involved in survival [68]. Access to antigen in the liver by T_RM_ cells might foster preferential antigen stimulation leading to the expression of pro-survival genes in these cells.

Although our results demonstrate that sporozoite-specific liver T_RM_ cells can provide highly efficient protection against infection, the preferential development of sporozoite specific T cell responses by RAS may have some drawbacks. Firstly, certain late antigens such as RPL6, which are highly protective [31], are not expanded as much upon RAS vaccination, and their full protective potential is therefore missed. This issue may be more prominent in human infections, where the liver stage lasts longer than in mice (7 days vs 2 days), and therefore T_RM_ cells targeting parasites during intrahepatic development may be comparably more relevant for protection. Secondly, reducing the generation of late-stage antigen-specific T cells may impair immunity against parasites with delayed development in the liver. Sporozoite-specific CD8^+^ T cells, such as those specific for CSP, are less efficient at protecting against *P. yoelii* than *P. berghei*, and this is associated to strongest late replication of the former parasite [24]. Moreover, unlike *P. berghei* RAS, repeated long interval *P. yoelii* RAS immunisation fails to improve protection in C57BL/6 mice [18]. Thirdly, we and others have observed that TRAP is highly variable across *P. falciparum* field isolates [31], and this is also true for CSP [88, 89]. Sporozoite surface antigens such as CSP or TRAP are directly exposed to immune attack and hence subjected to intense immune selection, which likely results in increased variability. Biasing the immune response towards highly polymorphic sporozoite antigens may increase the strain-specificity of RAS-mediated protection (or, similarly, of immunity from natural infection), reducing the capacity of memory T_RM_ cells generated to recognise and combat new infections. Administration of high doses of RAS, however, may aid initial expansion of protective T cells specific for less abundant sporozoite antigens and increase cross-strain immunity [25]. And fourthly, we have observed that the lifespan of liver T_RM_ cells generated by repeated RAS vaccination is strongly increased. This could potentially perpetuate the suppression of naïve responses to new antigens, not only further hindering the generation of T cell responses to liver stage antigens upon natural infection, but also to new sporozoite antigens from genetically diverse parasite haplotypes [90]. As we have seen here, those T cells are still able to respond to the new parasite, even when unable to exert protection. This could prevent the generation of immunity to novel epitopes, thereby reducing the breadth of immunity generated against newly encountered parasite strains. Together, these elements may establish a potent screen that prevents the generation of T_RM_ cell immunity of maximal efficacy against malaria parasite infection in the liver.

## Materials and Methods

### Mice, mosquitos, parasites and infections

All procedures were performed in strict accordance with the recommendations of the Australian code of practice for the care and use of animals for scientific purposes. The protocols were approved by the Melbourne Health Research Animal Ethics Committee, University of Melbourne (ethic project IDs: 1112347, 1814522, 20088).

Female C57BL/6 (B6), GFP [91], OT-I [46] and PbT-I [37] mice were used between 6-12 weeks of age and were bred and maintained at the Department of Microbiology and Immunology, The University of Melbourne. Animals used for the generation of the sporozoites were 4-5-week-old male Swiss Webster mice purchased from the Monash Animal Services (Melbourne, Victoria, Australia) and maintained at the School of Botany, The University of Melbourne, Australia. *Anopheles stephensi* mosquitoes (strain STE2/MRA-128 from The Malaria Research and Reference Reagent Resource Center) were reared and infected with *P. berghei* ANKA (*P. berghei*) as described [92]. *P. berghei* ANKA WT, *P. berghei* ANKA HsOVA [47] and *P. berghei* ANKA CS5M [43] sporozoites were dissected from mosquito salivary glands and resuspended in cold PBS. Freshly dissected *P. berghei* sporozoites were injected intravenously (i.v.) as indicated in the figure legends. Parasitemia was assessed by microscopic analysis of blood smears or by flow cytometry. Mice showing no evidence of blood-stage infection by day 11 after infection were considered sterilely protected. For blood stage infections, mice were injected i.v. with the indicated amount of *P. berghei* infected red blood cells (iRBC).

Heat killing of sporozoites was done by incubating freshly isolated sporozoites at 56°C for 45 minutes.

### Adoptive transfer of CD8^+^ T cells

PbT-I and OT-I CD8^+^ T cells were negatively enriched from the spleens and lymph nodes of mice from various genetic crosses as described [93]. 50,000 purified PbT-I cells in 0.2 mL PBS were injected i.v. into recipient mice. CellTrace^TM^ violet (CTV, Thermofisher) was used to coat PbT-I and OT-I cells following manufacturer’s instructions.

### Prime-and-trap vaccination

B6 mice were injected i.v. with the indicated doses of rat anti-Clec9A (clone 24/04-10B4) genetically fused to OVA (containing the OVA_257-264_ epitope) via a 4 Alanine linker to make the αClec9A-OVA mAb construct [94]. Recombinant adeno-associated virus (rAAV-OVA) was prepared and purified in house at the Centenary Institute or by the Vector and Genome Engineering Facility (at the Children Medical Research Institute, Sydney, Australia) over cesium chloride (CsCl)-density gradient centrifugation followed by dialysis as previously described (. This vector expresses a membrane bound form of OVA protein bicistronically with green fluorescent protein (GFP). αClec9A was injected with 5 nmol of a CpG oligonucleotide (CpG) generated by linking (5’ to 3’) CpG-2006 to CpG-21798 [95] (Integrated DNA Technologies, Coralville, IA, USA). For P&T vaccination, mice were injected i.v. with Clec9A mAb and the indicated vector gene copies (vgc) of rAAV-OVA on the same day.

### Organ processing for T cell analysis

Tissues were harvested from mice at different time points after immunization and finely chopped using curved scissors to generate single cell suspensions. For spleen cell preparations, red blood cells were lysed, and remaining cells were filtered through a 70 μm mesh. Liver cell suspensions were passed through a 70 μm mesh and resuspended in 35% isotonic Percoll. Cells were then centrifuged at 500g for 20 min at room temperature (RT), the pellet harvested, and then red cells lysed before further analysis.

### Flow cytometry

CD11a (2D7), CD8α (53-6.7) mAb were purchased from BD; CD44 (IM7), CD62L (MEL-14), CD69 (H1.2F3), from ThermoFisher Scientific (Waltham, MA, USA); CXCR3 (CXCR3-173), CXCR6 (SA05D1), CX3CR1 (SA011F11), from BioLegend (San Diego, CA, USA); H2-K^b^-PbRPL6_120-127_, H2-K^b^-PbRPA1_199-206_, H2-K^b^-OVA_257-264_ and H2-D^b^-PbTRAP_130-138_ tetramers were made in house. Dead cells were excluded by propidium iodide (PI) staining. For the analysis of memory CD8^+^ T cell populations in the spleen and the liver, tetramer^+^, PbT-I or OT-I CD8^+^ CD44^hi^ cells were subdivided into T_CM_, T_EM_ or T_RM_ based on CD69 and CD62L expression (T_CM_ CD62L^+^ CD69^-^, T_EM_ CD62L^-^ CD69^-^ and T_RM_ CD62L^-^CD69^+^, see Figure S1H). Parasitaemia was assessed by incubating ∼2μl tail blood with a 5 pg/mL Hoechst 33258 solution (ThermoFisher Scientific) in FACS buffer for 1 hour at 37°C. Parasites were discriminated from uninfected RBC using a 405 violet laser and a 450/50 filter. Cells were analyzed by flow cytometry on a FACS Canto, Fortessa or Fortessa X20 (BD Immunocytometry Systems, San Jose, CA, USA), using FACSDiva (BD Immunocytometry Systems) or FlowJo software (Tree Star, Ashland, OR, USA).

### Depletion of liver T cells

To deplete CXCR3^+^ cells, mice were intravenously injected with 2 doses (200 μg and then 100 μg) of anti-CXCR3 antibody (CXCR3-173, eBioscience) or Armenian Hamster IgG isotype control (eBio299Arm, eBioscience) 3 and 1 days before challenge with 200 live sporozoites [22]. To deplete CD8^+^ cells, mice were intravenously injected with 100 μg of anti-CD8 antibody (clone 2.43) or isotype control (GL117, IgG2a) one day before challenge.

### CD8 T cell tolerisation

Removal of TRAP specific T cells was achieved by injection of PbTRAP_130-138_ peptide in the absence of adjuvant [40, 44]. Mice received 3 intravenous doses of TRAP peptide diluted in PBS on days 7, 4 and 1 prior to the first dose of RAS, and then received additional injections a day before administration of subsequent doses of the vaccine. The first peptide dose was 300μg, and the rest were 100μg.

### Statistical analyses

Figures were generated using GraphPad Prism 10 (GraphPad Software, San Diego, CA, USA). Data are shown as mean values ± standard error of the mean (SEM). Statistical analyses were performed using GraphPad Prism 10. Unless otherwise stated, statistical comparisons of cell numbers in different groups were performed by log-transforming the data and using a Student’s t-test (2 groups) or one-way ANOVA followed by Tukey’s multiple comparisons test (>2 groups). Cell number values equal to 0 were converted to 1 to enable log transformation. *P*<0.05 was considered to indicate statistical significance. *, *P*<0.05; **, *P*<0.01; ***, *P*<0.001; ****, *P*<0.0001; n.s., not significant (*P*>0.05). Asterisks directly over groups denote statistical differences with the unvaccinated control group. Rates of sterile protection were compared using Fisher’s exact tests.

### Declaration of generative AI and AI-assisted technologies in the writing process

The authors used the AI-powered language models Perplexity AI and Microsoft Copilot for editorial suggestions to assist with improving the language and readability of this manuscript. The authors reviewed and edited the content as needed and take full responsibility for the content of the publication.

## Acknowledgements

We would like to thank Prof. Ian Cockburn for providing the CS5M parasites used in this work. We also thank Melanie Damtsis for technical assistance, the members of the W.R.H., L.K.M., S.M. and J.B. labs for discussions, the Doherty’s animal facility staff for mice husbandry and the Melbourne Brain Centre and ImmunoID Flow Cytometry Facilities for technical assistance.

## Author contributions

Conceptualization: D.F.R, W.R.H., L.B., M.N.dM.; Investigation: M.N.dM., D.F.R, Z.G.; Resources: S. G., G.I.M, A. C., M.H.L., P.B., I.C., K.Y.; Supervision: D.F.R, W.R.H., L.B.; Writing draft: D.F.R, W.R.H.; Writing -review and editing: M.N.dM., Z.G., S.G., M.H.L., K.Y., G.I.M., L.B.; Funding Acquisition: D.F.R., W.R.H.

## Data availability

All relevant data are within the manuscript and its supporting information files

## Supplementary figures

**Figure S1. Related to figure 1. CD8^+^ T cell depletion removes protection conferred by repeated RAS vaccination.** Mice vaccinated thrice with 10,000 RAS, one week apart, were treated with anti-CD8 antibodies on day 27 after the third RAS immunisation and were challenged with 200 live *P. berghei* sporozoites on day 30. (**A**) Rates of sterile protection. Numbers above columns denote numbers of protected mice / total numbers of mice per group. (**B**) Parasitemia at day 7 post-challenge. (**C**) Mice were bled on day 29 after the third RAS vaccination (i.e. 2 days after αCD8 treatment) and percentages of CD4^+^ and CD8^+^ T cells (showed as numbers above the CD8^+^ T cell gate) were measured in the blood using flow cytometry to verify CD8^+^ T cell depletion. An example of CD8^+^ T cell percentages in an untreated (top) and a treated (bottom) mouse are shown. One experiment was performed. Comparisons of sterile protection rates were done using Fisher’s exact tests. Parasitemia data were log-transformed and compared using one-way ANOVA and Tukey’s multiple comparisons tests. (**D-H)** Related to figure 1C, D. Distribution of memory T cell populations in the liver (**D, F)** and the spleen (**E, G)** in mice treated with αCXCR3 or isotype control mAb. **D** and **E** show total memory CD8^+^ T cells, whereas **F** and **G** show PbTRAP_130-138_ tetramer-specific memory CD8^+^ T cells. Cell numbers were log-transformed and compared using unpaired t-tests. Dark purple, pale purple and grey stats over the columns denote comparisons of T_RM_, T_EM_ and T_CM_ numbers respectively. (**H)** Representative gating strategy of liver cells, including lymphocytes, single cells, live CD8^+^ T cells (CD8^+^ Propidium Iodide [PI]^-^), memory T cells (CD44^high^), TRAP-tetramer^+^ cells (gated from memory T cells) and total (middle row) or TRAP-specific (bottom row) memory T cell subsets T_CM_ CD62L^+^ CD69^-^, T_EM_ CD62L^-^ CD69^-^ and T_RM_ CD62L^-^ CD69^+^. Panels titled “Isotype” and “αCXCR3” show total (middle row) or TRAP-specific (bottom row) memory T cells in the livers of a representative, isotype-treated and αCXCR3-treated mouse respectively. Numbers beside gates represent percentages over total cells in the plot.

**Figure S2. Related to figure 2. Abundance of liver CD8 T_RM_ cells specific for known *Plasmodium* antigens in mice vaccinated multiple times with WT RAS. A.** Expression of TRAP, RPL6 and RPA1 proteins in salivary gland sporozoites (sgSpz), injected sporozoites (bbSpz), exo-erythrocytic forms (EEF), merozoite and ring forms of *P. berghei*, as per the Malaria Cell Atlas [30]. **B.** Related to Figure 2A-C. Detailed distribution of memory CD8^+^ T cells of known (tetramer-positive, as indicated) or unknown (tetramer-negative) specificities in the spleen and the liver. Data were compared using one-way ANOVA and Tukey’s multiple comparisons test. The statistical analysis performed on liver data (dark asterisks) compared numbers of T_RM_ cells, and that in spleen data (pale asterisks) compared numbers of T_EM_ cells.

**Figure S3. Related to figure 2E-H. Detailed distribution of memory CD8^+^ T cells of known specificities in the spleen in mice vaccinated with 1x CS5M RAS or 3x CS5M RAS.** Data were compared using one-way ANOVA and Tukey’s multiple comparisons test. The statistical analysis performed on liver data (dark asterisks) compared numbers of T_RM_ cells.

**Figure S4. Related to figure 4. Tolerisation of TRAP-specific CD8^+^ T cells in RAS vaccinated mice. A.** Related to figure 4B-D. Representative flow cytometry charts showing depletion efficacy of TRAP specific cells. **B.** Related to figure 4C. Comparison of the parasitemias of those mice that were not sterilely protected.

**Figure S5**. **Related to figure 5. Distribution of TCR transgenic and endogenous T cells in RAS vaccinated mice. A.** Related to figure 5D-G. Distribution of PbT-I and endogenous, TRAP-, RPL6- and RPA1-specific memory T cells in the spleen and the liver as indicated. Numbers of T_EM_ cells were statistically compared in the spleen (pale green or purple asterisks), and numbers of T_RM_ cells were compared in the liver (dark asterisks). PbT-I and TRAP specific cell data were pooled from two independent experiments, and RPL6 and F4 cell data come from one experiment. Data were log-transformed and compared using one-way ANOVA and Tukey’s multiple comparisons test. **B.** Memory PbT-I cells in the liver on day 30 after transfer of 50,000 naïve PbT-I cells into mice that had been vaccinated with 10,000 RAS 6 days earlier (RAS + PbT-I), or one day later (PbT-I+RAS). Data were pooled from two independent experiments. **C.** Number of PbT-I cells, TRAP- and RPL6-specific CD8^+^ T cells in the blood on day 7 after the last RAS vaccination. Data come from one experiment and were log-transformed and compared using one-way ANOVA and Tukey’s multiple comparisons test. **D-F.** Related to figure 5H, I. **D.** Expression of Hsp70 across different life stages of the parasite, as per the Malaria Cell Atlas [30]. sgSpz, salivary gland sporozoites; bbSp, injected sporozoites; EEF, exo-erythrocytic forms. **E.** OT-I and PbT-I cell expansion after HsOVA HKS injection. Mice received 5x10^5^ naïve CellTrace Violet-coated OT-I and PbT-I cells one day before injection of 5.2-8x10^4^ HsOVA HKS, and numbers of divided OT-I and PbT-I cells were quantified in the spleen 4 days later. Data were pooled from two independent experiments, log-transformed and analysed using unpaired Student’s T-tests. **F.** Distribution of memory T cells of the indicated specificities in the liver and the spleen. Data were pooled from two independent experiments, log-transformed and analysed using two-way ANOVA and Tukey’s multiple comparisons test.

**Figure S6. Related to figure 5. Distribution of TCR transgenic and endogenous T cells in CS5M RAS vaccinated mice. A.** Expression of CSP across different life stages of the parasite, as per the Malaria Cell Atlas [30]. **B.** OT-I and PbT-I cell expansion after CS5M HKS injection. Mice received 5x10^5^ naïve CellTrace Violet-coated OT-I and PbT-I cells one day before injection of 4.2-4.5x10^4^ CS5M HKS, and numbers of divided OT-I and PbT-I cells were quantified in the spleen 6 days later. Data were pooled from two independent experiments, log-transformed and analysed using unpaired Student’s T tests. **C-D.** Related to figure 5I, J. Distribution of OVA-specific endogenous memory T cells (**C**) and other specificities as indicated (**D**) in the liver and the spleen. Data were pooled from two independent experiments, log-transformed and analysed using one-way ANOVA and Tukey’s multiple comparisons test. The statistical analysis performed on liver data compared numbers of T_RM_ cells

**Figure S7. Related to figure 5. Distribution of TCR transgenic and endogenous T cells in mice vaccinated with WT and CS5M RAS.** Memory CD8^+^ T cells were enumerated in the liver and the spleen of mice vaccinated with 2 doses of 5x10^3^ and 10x10^3^ RAS 4 days apart, then transferred with 50x10^3^ naïve OT-I cells and given a final dose of 5.1x10^3^ CS5M RAS 8 days later, or control mice receiving OT-I cells and one dose of 5.1x10^3^ CS5M RAS. Mice were euthanised on day 62 after the last sporozoite injection. Data were generated in one experiment, log-transformed and analysed using one-way ANOVA and Tukey’s multiple comparisons test. The statistical analysis performed on liver data compared numbers of T_RM_ cells.

**Figure S8. Related to figure 5K, L. Distribution of TCR transgenic and endogenous memory T cells in RAS-vaccinated mice that were efficiently or suboptimally tolerised for PbTRAP_130-138_. A.** Memory PbT-I cells in the spleen. **B.** TRAP-specific memory CD8^+^ T cells in the spleen. **C-D**. Numbers of TRAP-specific (**C**) and PbT-I (**D**) memory cells in the liver as in figure 5K, L, but mice in the 3xRAS group in which TRAP tolerisation worked efficiently (effT) or suboptimally (subT) were separated into different columns. **E.** Endogenous memory CD8^+^ T cells of undefined specificities (non-TRAP) in the liver. **F.** Endogenous memory CD8^+^ T cells of undefined specificities (non-TRAP) in the spleen.

## References

1. Geneva: World Health Organization W. World malaria report 2023. Licence: CC BY-NC-SA 3.0 IGO2023.

2. Bloom DE. The Value of Vaccination. Springer New York; 2011. p. 1-8.

3. Nussenzweig RS, Vanderberg J, Most H, Orton C. Protective Immunity produced by the Injection of X-irradiated Sporozoites of Plasmodium berghei. Nature. 1967;216(5111):160-2. doi: 10.1038/216160a0.

4. Gwadz RW, Cochrane AH, Nussenzweig V, Nussenzweig RS. Preliminary studies on vaccination of rhesus monkeys with irradiated sporozoites of Plasmodium knowlesi and characterization of surface antigens of these parasites. Bull World Health Organ. 1979;57 Suppl 1(Suppl):165-73. PubMed PMID: 120766; PubMed Central PMCID: PMCPMC2395714.

5. Clyde DF, Most H, McCarthy VC, Vanderberg JP. Immunization of man against sporozite-induced falciparum malaria. Am J Med Sci. 1973;266(3):169–77. doi: 10.1097/00000441-197309000-00002. PubMed PMID: 4583408.

6. Mellouk S, Lunel F, Sedegah M, Beaudoin RL, Druilhe P. Protection against malaria induced by irradiated sporozoites. Lancet. 1990;335(8691):721. doi: 10.1016/0140-6736(90)90832-p. PubMed PMID: 1969073.

7. Scheller LF, Stump KC, Azad AF. Plasmodium berghei: production and quantitation of hepatic stages derived from irradiated sporozoites in rats and mice. J Parasitol. 1995;81(1):58–62. PubMed PMID: 7876979.

8. Silvie O, Semblat JP, Franetich JF, Hannoun L, Eling W, Mazier D. Effects of irradiation on Plasmodium falciparum sporozoite hepatic development: implications for the design of pre-erythrocytic malaria vaccines. Parasite Immunol. 2002;24(4):221-3. Epub 2002/05/16. doi: 10.1046/j.1365-3024.2002.00450.x. PubMed PMID: 12010486.

9. Seder RA, Chang LJ, Enama ME, Zephir KL, Sarwar UN, Gordon IJ, et al. Protection against malaria by intravenous immunization with a nonreplicating sporozoite vaccine. Science. 2013;341(6152):1359-65. Epub 2013/08/10. doi: 10.1126/science.1241800. PubMed PMID: 23929949.

10. Schofield L, Villaquiran J, Ferreira A, Schellekens H, Nussenzweig R, Nussenzweig V. γ Interferon, CD8+ T cells and antibodies required for immunity to malaria sporozoites. Nature. 1987;330(6149):664-6. doi: 10.1038/330664a0.

11. Oliveira GA, Kumar KA, Calvo-Calle JM, Othoro C, Altszuler D, Nussenzweig V, et al. Class II-restricted protective immunity induced by malaria sporozoites. Infect Immun. 2008;76(3):1200–6. Epub 2007/12/28. doi: 10.1128/IAI.00566-07. PubMed PMID: 18160479; PubMed Central PMCID: PMCPMC2258813.

12. Schmidt NW, Butler NS, Harty JT. CD8 T cell immunity to Plasmodium permits generation of protective antibodies after repeated sporozoite challenge. Vaccine. 2009;27(44):6103–6. Epub 2009/08/29. doi: 10.1016/j.vaccine.2009.08.025. PubMed PMID: 19712771; PubMed Central PMCID: PMCPMC2759845.

13. Ishizuka AS, Lyke KE, DeZure A, Berry AA, Richie TL, Mendoza FH, et al. Protection against malaria at 1 year and immune correlates following PfSPZ vaccination. Nat Med. 2016;22(6):614–23. Epub 2016/05/10. doi: 10.1038/nm.4110. PubMed PMID: 27158907.

14. Weiss WR, Sedegah M, Beaudoin RL, Miller LH, Good MF. CD8+ T cells (cytotoxic/suppressors) are required for protection in mice immunized with malaria sporozoites. Proc Natl Acad Sci U S A. 1988;85(2):573–6. Epub 1988/01/01. doi: 10.1073/pnas.85.2.573. PubMed PMID: 2963334; PubMed Central PMCID: PMCPMC279593.

15. Doolan DL, Hoffman SL. The complexity of protective immunity against liver-stage malaria. J Immunol. 2000;165(3):1453–62. doi: 10.4049/jimmunol.165.3.1453. PubMed PMID: 10903750.

16. Weiss WR, Jiang CG. Protective CD8+ T lymphocytes in primates immunized with malaria sporozoites. PLoS One. 2012;7(2):e31247. Epub 2012/02/23. doi: 10.1371/journal.pone.0031247. PubMed PMID: 22355349; PubMed Central PMCID: PMCPMC3280278.

17. Guebre-Xabier M, Schwenk R, Krzych U. Memory phenotype CD8+ T cells persist in livers of mice protected against malaria by immunization with attenuatedPlasmodium bergheisporozoites. European Journal of Immunology. 1999;29(12):3978–86. doi: 10.1002/(sici)1521-4141(199912)29:12<3978::aid-immu3978>3.0.co;2-0.

18. Schmidt NW, Butler NS, Badovinac VP, Harty JT. Extreme CD8 T cell requirements for anti-malarial liver-stage immunity following immunization with radiation attenuated sporozoites. PLoS Pathog. 2010;6(7):e1000998. Epub 2010/07/27. doi: 10.1371/journal.ppat.1000998. PubMed PMID: 20657824; PubMed Central PMCID: PMCPMC2904779.

19. Epstein JE, Tewari K, Lyke KE, Sim BK, Billingsley PF, Laurens MB, et al. Live attenuated malaria vaccine designed to protect through hepatic CD8(+) T cell immunity. Science. 2011;334(6055):475-80. Epub 2011/09/10. doi: 10.1126/science.1211548. PubMed PMID: 21903775.

20. Fernandez-Ruiz D, de Menezes MN, Holz LE, Ghilas S, Heath WR, Beattie L. Harnessing liver-resident memory T cells for protection against malaria. Expert Rev Vaccines. 2021;20(2):127–41. Epub 22/01/2021. doi: 10.1080/14760584.2021.1881485. PubMed PMID: 33501877.

21. Lefebvre MN, Surette FA, Anthony SM, Vijay R, Jensen IJ, Pewe LL, et al. Expeditious recruitment of circulating memory CD8 T cells to the liver facilitates control of malaria. Cell Rep. 2021;37(5):109956. doi: 10.1016/j.celrep.2021.109956. PubMed PMID: 34731605; PubMed Central PMCID: PMCPMC8628427.

22. Fernandez-Ruiz D, Ng WY, Holz LE, Ma JZ, Zaid A, Wong YC, et al. Liver-Resident Memory CD8(+) T Cells Form a Front-Line Defense against Malaria Liver-Stage Infection. Immunity. 2016;45(4):889–902. Epub 2016/10/21. doi: 10.1016/j.immuni.2016.08.011. PubMed PMID: 27692609.

23. Jaffe RI, Lowell GH, Gordon DM. Differences in Susceptibility Among Mouse Strains to Infection with Plasmodium Berghei (Anka Clone) Sporozoites and its Relationship to Protection by Gamma-Irradiated Sporozoites. The American Journal of Tropical Medicine and Hygiene. 1990;42(4):309–13. doi: 10.4269/ajtmh.1990.42.309.

24. Schmidt NW, Butler NS, Harty JT. Plasmodium-host interactions directly influence the threshold of memory CD8 T cells required for protective immunity. J Immunol. 2011;186(10):5873–84. Epub 2011/04/05. doi: 10.4049/jimmunol.1100194. PubMed PMID: 21460205; PubMed Central PMCID: PMCPMC3087867.

25. Lyke KE, Ishizuka AS, Berry AA, Chakravarty S, DeZure A, Enama ME, et al. Attenuated PfSPZ Vaccine induces strain-transcending T cells and durable protection against heterologous controlled human malaria infection. Proc Natl Acad Sci U S A. 2017;114(10):2711–6. Epub 2017/02/23. doi: 10.1073/pnas.1615324114. PubMed PMID: 28223498; PubMed Central PMCID: PMCPMC5347610.

26. Landau I, Boulard Y. Life cycles and morphology. In: Killick-Kendrick R, Peters W, editors. Rodent Malaria. London: Academic Press Inc.; 1978. p. 53-84.

27. Sturm A, Amino R, van de Sand C, Regen T, Retzlaff S, Rennenberg A, et al. Manipulation of host hepatocytes by the malaria parasite for delivery into liver sinusoids. Science. 2006;313(5791):1287-90. Epub 2006/08/05. doi: 10.1126/science.1129720. PubMed PMID: 16888102.

28. Vaughan AM, Mikolajczak SA, Wilson EM, Grompe M, Kaushansky A, Camargo N, et al. Complete Plasmodium falciparum liver-stage development in liver-chimeric mice. J Clin Invest. 2012;122(10):3618–28. Epub 2012/09/22. doi: 10.1172/JCI62684. PubMed PMID: 22996664; PubMed Central PMCID: PMCPMC3461911.

29. Reid AJ, Talman AM, Bennett HM, Gomes AR, Sanders MJ, Illingworth CJR, et al. Single-cell RNA-seq reveals hidden transcriptional variation in malaria parasites. Elife. 2018;7. Epub 20180327. doi: 10.7554/eLife.33105. PubMed PMID: 29580379; PubMed Central PMCID: PMCPMC5871331.

30. Howick VM, Russell AJC, Andrews T, Heaton H, Reid AJ, Natarajan K, et al. The Malaria Cell Atlas: Single parasite transcriptomes across the complete Plasmodium life cycle. Science. 2019;365(6455). Epub 2019/08/24. doi: 10.1126/science.aaw2619. PubMed PMID: 31439762; PubMed Central PMCID: PMCPMC7056351.

31. Valencia-Hernandez AM, Ng WY, Ghazanfari N, Ghilas S, de Menezes MN, Holz LE, et al. A Natural Peptide Antigen within the Plasmodium Ribosomal Protein RPL6 Confers Liver TRM Cell-Mediated Immunity against Malaria in Mice. Cell Host Microbe. 2020;27(6):950–62 e7. Epub 2020/05/13. doi: 10.1016/j.chom.2020.04.010. PubMed PMID: 32396839.

32. Amino R, Thiberge S, Martin B, Celli S, Shorte S, Frischknecht F, et al. Quantitative imaging of Plasmodium transmission from mosquito to mammal. Nat Med. 2006;12(2):220–4. Epub 2006/01/24. doi: 10.1038/nm1350. PubMed PMID: 16429144.

33. Butler NS, Schmidt NW, Vaughan AM, Aly AS, Kappe SH, Harty JT. Superior antimalarial immunity after vaccination with late liver stage-arresting genetically attenuated parasites. Cell Host Microbe. 2011;9(6):451–62. Epub 2011/06/15. doi: 10.1016/j.chom.2011.05.008. PubMed PMID: 21669394; PubMed Central PMCID: PMCPMC3117254.

34. Yewdell JW. Confronting Complexity: Real-World Immunodominance in Antiviral CD8+ T Cell Responses. Immunity. 2006;25(4):533–43. doi: 10.1016/j.immuni.2006.09.005.

35. Jenkins MK, Moon JJ. The role of naive T cell precursor frequency and recruitment in dictating immune response magnitude. J Immunol. 2012;188(9):4135–40. Epub 2012/04/21. doi: 10.4049/jimmunol.1102661. PubMed PMID: 22517866; PubMed Central PMCID: PMCPMC3334329.

36. Kumar KA, Sano G, Boscardin S, Nussenzweig RS, Nussenzweig MC, Zavala F, et al. The circumsporozoite protein is an immunodominant protective antigen in irradiated sporozoites. Nature. 2006;444(7121):937-40. Epub 2006/12/08. doi: 10.1038/nature05361. PubMed PMID: 17151604.

37. Lau LS, Fernandez-Ruiz D, Mollard V, Sturm A, Neller MA, Cozijnsen A, et al. CD8+ T cells from a novel T cell receptor transgenic mouse induce liver-stage immunity that can be boosted by blood-stage infection in rodent malaria. PLoS Pathog. 2014;10(5):e1004135. Epub 2014/05/24. doi: 10.1371/journal.ppat.1004135. PubMed PMID: 24854165; PubMed Central PMCID: PMCPMC4031232.

38. Badovinac VP, Haring JS, Harty JT. Initial T cell receptor transgenic cell precursor frequency dictates critical aspects of the CD8(+) T cell response to infection. Immunity. 2007;26(6):827–41. Epub 2007/06/09. doi: 10.1016/j.immuni.2007.04.013. PubMed PMID: 17555991; PubMed Central PMCID: PMCPMC1989155.

39. Nganou-Makamdop K, van Gemert GJ, Arens T, Hermsen CC, Sauerwein RW. Long term protection after immunization with P. berghei sporozoites correlates with sustained IFNgamma responses of hepatic CD8+ memory T cells. PLoS One. 2012;7(5):e36508. Epub 2012/05/09. doi: 10.1371/journal.pone.0036508. PubMed PMID: 22563506; PubMed Central PMCID: PMCPMC3341355.

40. Hafalla JC, Bauza K, Friesen J, Gonzalez-Aseguinolaza G, Hill AV, Matuschewski K. Identification of targets of CD8(+) T cell responses to malaria liver stages by genome-wide epitope profiling. PLoS Pathog. 2013;9(5):e1003303. Epub 2013/05/16. doi: 10.1371/journal.ppat.1003303. PubMed PMID: 23675294; PubMed Central PMCID: PMCPMC3649980.

41. Hall N, Karras M, Raine JD, Carlton JM, Kooij TW, Berriman M, et al. A comprehensive survey of the Plasmodium life cycle by genomic, transcriptomic, and proteomic analyses. Science. 2005;307(5706):82-6. Epub 2005/01/08. doi: 10.1126/science.1103717. PubMed PMID: 15637271.

42. Lau LS, Fernandez-Ruiz D, Davey GM, de Koning-Ward TF, Papenfuss AT, Carbone FR, et al. Blood-stage Plasmodium berghei infection generates a potent, specific CD8+ T-cell response despite residence largely in cells lacking MHC I processing machinery. J Infect Dis. 2011;204(12):1989–96. Epub 2011/10/15. doi: 10.1093/infdis/jir656. PubMed PMID: 21998471.

43. Cockburn IA, Tse SW, Radtke AJ, Srinivasan P, Chen YC, Sinnis P, et al. Dendritic cells and hepatocytes use distinct pathways to process protective antigen from plasmodium in vivo. PLoS Pathog. 2011;7(3):e1001318. Epub 2011/03/30. doi: 10.1371/journal.ppat.1001318. PubMed PMID: 21445239; PubMed Central PMCID: PMCPMC3060173.

44. Redmond WL, Marincek BC, Sherman LA. Distinct requirements for deletion versus anergy during CD8 T cell peripheral tolerance in vivo. J Immunol. 2005;174(4):2046–53. doi: 10.4049/jimmunol.174.4.2046. PubMed PMID: 15699134.

45. Murphy SC, Kas A, Stone BC, Bevan MJ. A T-cell response to a liver-stage Plasmodium antigen is not boosted by repeated sporozoite immunizations. Proc Natl Acad Sci U S A. 2013;110(15):6055–60. Epub 2013/03/27. doi: 10.1073/pnas.1303834110. PubMed PMID: 23530242; PubMed Central PMCID: PMCPMC3625320.

46. Hogquist KA, Jameson SC, Heath WR, Howard JL, Bevan MJ, Carbone FR. T cell receptor antagonist peptides induce positive selection. Cell. 1994;76(1):17–27. doi: 10.1016/0092-8674(94)90169-4.

47. Miyakoda M, Kimura D, Yuda M, Chinzei Y, Shibata Y, Honma K, et al. Malaria-specific and nonspecific activation of CD8+ T cells during blood stage of Plasmodium berghei infection. J Immunol. 2008;181(2):1420–8. doi: 10.4049/jimmunol.181.2.1420. PubMed PMID: 18606696.

48. Rios KT, McGee JP, Sebastian A, Moritz RL, Feric M, Absalon S, et al. Global Release of Translational Repression Across Plasmodium’s Host-to-Vector Transmission Event. bioRxiv. 2024. Epub 20240316. doi: 10.1101/2024.02.01.577866. PubMed PMID: 38352447; PubMed Central PMCID: PMCPMC10862809.

49. Ghilas S, Enders MH, May R, Holz LE, Fernandez-Ruiz D, Cozijnsen A, et al. Development of Plasmodium-specific liver-resident memory CD8(+) T cells after heat-killed sporozoite immunization in mice. Eur J Immunol. 2021;51(5):1153–65. Epub 23/01/2021. doi: 10.1002/eji.202048757. PubMed PMID: 33486759.

50. Kimura K, Kimura D, Matsushima Y, Miyakoda M, Honma K, Yuda M, et al. CD8+ T cells specific for a malaria cytoplasmic antigen form clusters around infected hepatocytes and are protective at the liver stage of infection. Infect Immun. 2013;81(10):3825–34. Epub 2013/07/31. doi: 10.1128/IAI.00570-13. PubMed PMID: 23897612; PubMed Central PMCID: PMCPMC3811763.

51. Zhou Y, Ramachandran V, Kumar KA, Westenberger S, Refour P, Zhou B, et al. Evidence-based annotation of the malaria parasite’s genome using comparative expression profiling. PLoS One. 2008;3(2):e1570. Epub 2008/02/14. doi: 10.1371/journal.pone.0001570. PubMed PMID: 18270564; PubMed Central PMCID: PMCPMC2215772.

52. Jongo SA, Church LWP, Mtoro AT, Schindler T, Chakravarty S, Ruben AJ, et al. Increase of Dose Associated With Decrease in Protection Against Controlled Human Malaria Infection by PfSPZ Vaccine in Tanzanian Adults. Clin Infect Dis. 2020;71(11):2849–57. Epub 2019/11/30. doi: 10.1093/cid/ciz1152. PubMed PMID: 31782768; PubMed Central PMCID: PMCPMC7947995.

53. Steinert EM, Schenkel JM, Fraser KA, Beura LK, Manlove LS, Igyarto BZ, et al. Quantifying Memory CD8 T Cells Reveals Regionalization of Immunosurveillance. Cell. 2015;161(4):737–49. Epub 2015/05/11. doi: 10.1016/j.cell.2015.03.031. PubMed PMID: 25957682; PubMed Central PMCID: PMCPMC4426972.

54. Kumar BV, Ma W, Miron M, Granot T, Guyer RS, Carpenter DJ, et al. Human Tissue-Resident Memory T Cells Are Defined by Core Transcriptional and Functional Signatures in Lymphoid and Mucosal Sites. Cell Rep. 2017;20(12):2921–34. Epub 2017/09/21. doi: 10.1016/j.celrep.2017.08.078. PubMed PMID: 28930685; PubMed Central PMCID: PMCPMC5646692.

55. Steinbach K, Vincenti I, Kreutzfeldt M, Page N, Muschaweckh A, Wagner I, et al. Brain-resident memory T cells represent an autonomous cytotoxic barrier to viral infection. J Exp Med. 2016;213(8):1571–87. Epub 2016/07/06. doi: 10.1084/jem.20151916. PubMed PMID: 27377586; PubMed Central PMCID: PMCPMC4986533.

56. Hombrink P, Helbig C, Backer RA, Piet B, Oja AE, Stark R, et al. Programs for the persistence, vigilance and control of human CD8(+) lung-resident memory T cells. Nat Immunol. 2016;17(12):1467–78. Epub 2016/11/01. doi: 10.1038/ni.3589. PubMed PMID: 27776108.

57. Oja AE, Piet B, Helbig C, Stark R, van der Zwan D, Blaauwgeers H, et al. Trigger-happy resident memory CD4(+) T cells inhabit the human lungs. Mucosal Immunol. 2018;11(3):654–67. Epub 2017/11/16. doi: 10.1038/mi.2017.94. PubMed PMID: 29139478.

58. Schenkel JM, Fraser KA, Beura LK, Pauken KE, Vezys V, Masopust D. T cell memory. Resident memory CD8 T cells trigger protective innate and adaptive immune responses. Science. 2014;346(6205):98-101. Epub 2014/08/30. doi: 10.1126/science.1254536. PubMed PMID: 25170049; PubMed Central PMCID: PMCPMC4449618.

59. Wu T, Hu Y, Lee YT, Bouchard KR, Benechet A, Khanna K, et al. Lung-resident memory CD8 T cells (TRM) are indispensable for optimal cross-protection against pulmonary virus infection. J Leukoc Biol. 2014;95(2):215–24. Epub 2013/09/06. doi: 10.1189/jlb.0313180. PubMed PMID: 24006506; PubMed Central PMCID: PMCPMC3896663.

60. Shin H, Iwasaki A. A vaccine strategy that protects against genital herpes by establishing local memory T cells. Nature. 2012;491(7424):463-7. Epub 2012/10/19. doi: 10.1038/nature11522. PubMed PMID: 23075848; PubMed Central PMCID: PMCPMC3499630.

61. Teijaro JR, Turner D, Pham Q, Wherry EJ, Lefrancois L, Farber DL. Cutting edge: Tissue-retentive lung memory CD4 T cells mediate optimal protection to respiratory virus infection. J Immunol. 2011;187(11):5510–4. Epub 2011/11/08. doi: 10.4049/jimmunol.1102243. PubMed PMID: 22058417; PubMed Central PMCID: PMCPMC3221837.

62. Gebhardt T, Wakim LM, Eidsmo L, Reading PC, Heath WR, Carbone FR. Memory T cells in nonlymphoid tissue that provide enhanced local immunity during infection with herpes simplex virus. Nat Immunol. 2009;10(5):524–30. Epub 2009/03/24. doi: 10.1038/ni.1718. PubMed PMID: 19305395.

63. Ganley M, Holz LE, Minnell JJ, de Menezes MN, Burn OK, Poa KCY, et al. mRNA vaccine against malaria tailored for liver-resident memory T cells. Nat Immunol. 2023;24(9):1487–98. Epub 20230720. doi: 10.1038/s41590-023-01562-6. PubMed PMID: 37474653.

64. Holz LE, Chua YC, de Menezes MN, Anderson RJ, Draper SL, Compton BJ, et al. Glycolipid-peptide vaccination induces liver-resident memory CD8(+) T cells that protect against rodent malaria. Sci Immunol. 2020;5(48):eaaz8035. Epub 2020/06/28. doi: 10.1126/sciimmunol.aaz8035. PubMed PMID: 32591409.

65. Park SL, Zaid A, Hor JL, Christo SN, Prier JE, Davies B, et al. Local proliferation maintains a stable pool of tissue-resident memory T cells after antiviral recall responses. Nat Immunol. 2018;19(2):183–91. Epub 2018/01/10. doi: 10.1038/s41590-017-0027-5. PubMed PMID: 29311695.

66. Beura LK, Wijeyesinghe S, Thompson EA, Macchietto MG, Rosato PC, Pierson MJ, et al. T Cells in Nonlymphoid Tissues Give Rise to Lymph-Node-Resident Memory T Cells. Immunity. 2018;48(2):327–38 e5. Epub 2018/02/22. doi: 10.1016/j.immuni.2018.01.015. PubMed PMID: 29466758; PubMed Central PMCID: PMCPMC5828517.

67. Slutter B, Van Braeckel-Budimir N, Abboud G, Varga SM, Salek-Ardakani S, Harty JT. Dynamics of influenza-induced lung-resident memory T cells underlie waning heterosubtypic immunity. Sci Immunol. 2017;2(7). Epub 2017/08/08. doi: 10.1126/sciimmunol.aag2031. PubMed PMID: 28783666; PubMed Central PMCID: PMCPMC5590757.

68. Van Braeckel-Budimir N, Varga SM, Badovinac VP, Harty JT. Repeated Antigen Exposure Extends the Durability of Influenza-Specific Lung-Resident Memory CD8(+) T Cells and Heterosubtypic Immunity. Cell Rep. 2018;24(13):3374–82 e3. Epub 2018/09/27. doi: 10.1016/j.celrep.2018.08.073. PubMed PMID: 30257199; PubMed Central PMCID: PMCPMC6258017.

69. Wirth TC, Xue H-H, Rai D, Sabel JT, Bair T, Harty JT, et al. Repetitive Antigen Stimulation Induces Stepwise Transcriptome Diversification but Preserves a Core Signature of Memory CD8+ T Cell Differentiation. Immunity. 2010;33(1):128–40. doi: 10.1016/j.immuni.2010.06.014.

70. Tarun AS, Peng X, Dumpit RF, Ogata Y, Silva-Rivera H, Camargo N, et al. A combined transcriptome and proteome survey of malaria parasite liver stages. Proc Natl Acad Sci U S A. 2008;105(1):305–10. Epub 2008/01/04. doi: 10.1073/pnas.0710780104. PubMed PMID: 18172196; PubMed Central PMCID: PMCPMC2224207.

71. Weiss WR, Mellouk S, Houghten RA, Sedegah M, Kumar S, Good MF, et al. Cytotoxic T cells recognize a peptide from the circumsporozoite protein on malaria-infected hepatocytes. The Journal of experimental medicine. 1990;171(3):763–73. doi: 10.1084/jem.171.3.763.

72. Rogers WO, Malik A, Mellouk S, Nakamura K, Rogers MD, Szarfman A, et al. Characterization of Plasmodium falciparum sporozoite surface protein 2. Proceedings of the National Academy of Sciences. 1992;89(19):9176–80. doi: 10.1073/pnas.89.19.9176.

73. Van Braeckel-Budimir N, Gras S, Ladell K, Josephs TM, Pewe L, Urban SL, et al. A T Cell Receptor Locus Harbors a Malaria-Specific Immune Response Gene. Immunity. 2017;47(5):835–47 e4. Epub 2017/11/19. doi: 10.1016/j.immuni.2017.10.013. PubMed PMID: 29150238; PubMed Central PMCID: PMCPMC5724374.

74. Chakravarty S, Cockburn IA, Kuk S, Overstreet MG, Sacci JB, Zavala F. CD8+ T lymphocytes protective against malaria liver stages are primed in skin-draining lymph nodes. Nat Med. 2007;13(9):1035–41. Epub 2007/08/21. doi: 10.1038/nm1628. PubMed PMID: 17704784.

75. Wen-yue X, Xing-xiang W, Jie Q, Jian-hua D, Fu-sheng H. Plasmodium yoelii: influence of immune modulators on the development of the liver stage. Exp Parasitol. 2010;126(2):254–8. Epub 20100521. doi: 10.1016/j.exppara.2010.05.005. PubMed PMID: 20493849.

76. Coppi A, Pinzon-Ortiz C, Hutter C, Sinnis P. The Plasmodium circumsporozoite protein is proteolytically processed during cell invasion. J Exp Med. 2005;201(1):27–33. doi: 10.1084/jem.20040989. PubMed PMID: 15630135; PubMed Central PMCID: PMCPMC1995445.

77. Silvie O, Franetich J-F, Charrin S, Mueller MS, Siau A, Bodescot M, et al. A Role for Apical Membrane Antigen 1 during Invasion of Hepatocytes by Plasmodium falciparum Sporozoites. Journal of Biological Chemistry. 2004;279(10):9490–6. doi: 10.1074/jbc.m311331200.

78. Kaushansky A, Metzger PG, Douglass AN, Mikolajczak SA, Lakshmanan V, Kain HS, et al. Malaria parasite liver stages render host hepatocytes susceptible to mitochondria-initiated apoptosis. Cell Death & Disease. 2013;4(8):e762-e. doi: 10.1038/cddis.2013.286.

79. Kaech SM, Ahmed R. Memory CD8+ T cell differentiation: initial antigen encounter triggers a developmental program in naive cells. Nat Immunol. 2001;2(5):415–22. doi: 10.1038/87720. PubMed PMID: 11323695; PubMed Central PMCID: PMCPMC3760150.

80. Mitchell DM, Ravkov EV, Williams MA. Distinct Roles for IL-2 and IL-15 in the Differentiation and Survival of CD8+ Effector and Memory T Cells. The Journal of Immunology. 2010;184(12):6719–30. doi: 10.4049/jimmunol.0904089.

81. Hafalla JC, Sano G, Carvalho LH, Morrot A, Zavala F. Short-term antigen presentation and single clonal burst limit the magnitude of the CD8(+) T cell responses to malaria liver stages. Proc Natl Acad Sci U S A. 2002;99(18):11819–24. Epub 2002/08/20. doi: 10.1073/pnas.182189999. PubMed PMID: 12185251; PubMed Central PMCID: PMCPMC129352.

82. Hafalla JC, Morrot A, Sano G, Milon G, Lafaille JJ, Zavala F. Early self-regulatory mechanisms control the magnitude of CD8+ T cell responses against liver stages of murine malaria. J Immunol. 2003;171(2):964–70. doi: 10.4049/jimmunol.171.2.964. PubMed PMID: 12847268.

83. Romero P, Maryanski JL, Corradin G, Nussenzweig RS, Nussenzweig V, Zavala F. Cloned cytotoxic T cells recognize an epitope in the circumsporozoite protein and protect against malaria. Nature. 1989;341(6240):323-6. doi: 10.1038/341323a0.

84. Gruner AC, Mauduit M, Tewari R, Romero JF, Depinay N, Kayibanda M, et al. Sterile protection against malaria is independent of immune responses to the circumsporozoite protein. PLoS One. 2007;2(12):e1371. Epub 2007/12/27. doi: 10.1371/journal.pone.0001371. PubMed PMID: 18159254; PubMed Central PMCID: PMCPMC2147056.

85. Muller K, Gibbins MP, Roberts M, Reyes-Sandoval A, Hill AVS, Draper SJ, et al. Low immunogenicity of malaria pre-erythrocytic stages can be overcome by vaccination. EMBO Mol Med. 2021;13(4):e13390. Epub 2021/03/13. doi: 10.15252/emmm.202013390. PubMed PMID: 33709544; PubMed Central PMCID: PMCPMC8033512.

86. Doll KL, Pewe LL, Kurup SP, Harty JT. Discriminating Protective from Nonprotective Plasmodium-Specific CD8+ T Cell Responses. J Immunol. 2016;196(10):4253–62. Epub 2016/04/17. doi: 10.4049/jimmunol.1600155. PubMed PMID: 27084099; PubMed Central PMCID: PMCPMC4868661.

87. Rai D, Martin MD, Badovinac VP. The Longevity of Memory CD8 T Cell Responses after Repetitive Antigen Stimulations. The Journal of Immunology. 2014;192(12):5652–9. doi: 10.4049/jimmunol.1301063.

88. Kidgell C, Volkman SK, Daily J, Borevitz JO, Plouffe D, Zhou Y, et al. A Systematic Map of Genetic Variation in Plasmodium falciparum. PLoS Pathogens. 2006;2(6):e57. doi: 10.1371/journal.ppat.0020057.

89. Gandhi K, Thera MA, Coulibaly D, Traoré K, Guindo AB, Ouattara A, et al. Variation in the Circumsporozoite Protein of Plasmodium falciparum: Vaccine Development Implications. PLoS ONE. 2014;9(7):e101783. doi: 10.1371/journal.pone.0101783.

90. Naung MT, Martin E, Munro J, Mehra S, Guy AJ, Laman M, et al. Global diversity and balancing selection of 23 leading Plasmodium falciparum candidate vaccine antigens. PLOS Computational Biology. 2022;18(2):e1009801. doi: 10.1371/journal.pcbi.1009801.

91. Okabe M, Ikawa M, Kominami K, Nakanishi T, Nishimune Y. ‘Green mice’ as a source of ubiquitous green cells. FEBS Letters. 1997;407(3):313–9. doi: 10.1016/s0014-5793(97)00313-x.

92. Benedict MQ. Care and maintenance of anopheline mosquito colonies. In: Crampton JM, Beard CB, Louis C, editors. The Molecular Biology of Insect Disease Vectors: A Methods Manual. Dordrecht: Springer Netherlands; 1997. p. 3-12.

93. Smith CM, Belz GT, Wilson NS, Villadangos JA, Shortman K, Carbone FR, et al. Cutting edge: conventional CD8 alpha+ dendritic cells are preferentially involved in CTL priming after footpad infection with herpes simplex virus-1. J Immunol. 2003;170(9):4437–40. Epub 2003/04/23. doi: 10.4049/jimmunol.170.9.4437. PubMed PMID: 12707318.

94. Lahoud MH, Ahmet F, Kitsoulis S, Wan SS, Vremec D, Lee CN, et al. Targeting antigen to mouse dendritic cells via Clec9A induces potent CD4 T cell responses biased toward a follicular helper phenotype. J Immunol. 2011;187(2):842–50. Epub 2011/06/17. doi: 10.4049/jimmunol.1101176. PubMed PMID: 21677141.

95. Li J, Panetta F, O’Keeffe M, Leal Rojas IM, Radford KJ, Zhang J-G, et al. Elucidating the Motif for CpG Oligonucleotide Binding to the Dendritic Cell Receptor DEC-205 Leads to Improved Adjuvants for Liver-Resident Memory. The Journal of Immunology. 2021:ji2001153. doi: 10.4049/jimmunol.2001153.

