## Supplementary figures for "Long lived liver-resident memory T cells of biased specificities for abundant sporozoite antigens drive malaria protection by radiation-attenuated sporozoite vaccination"

### Supplementary figure 1

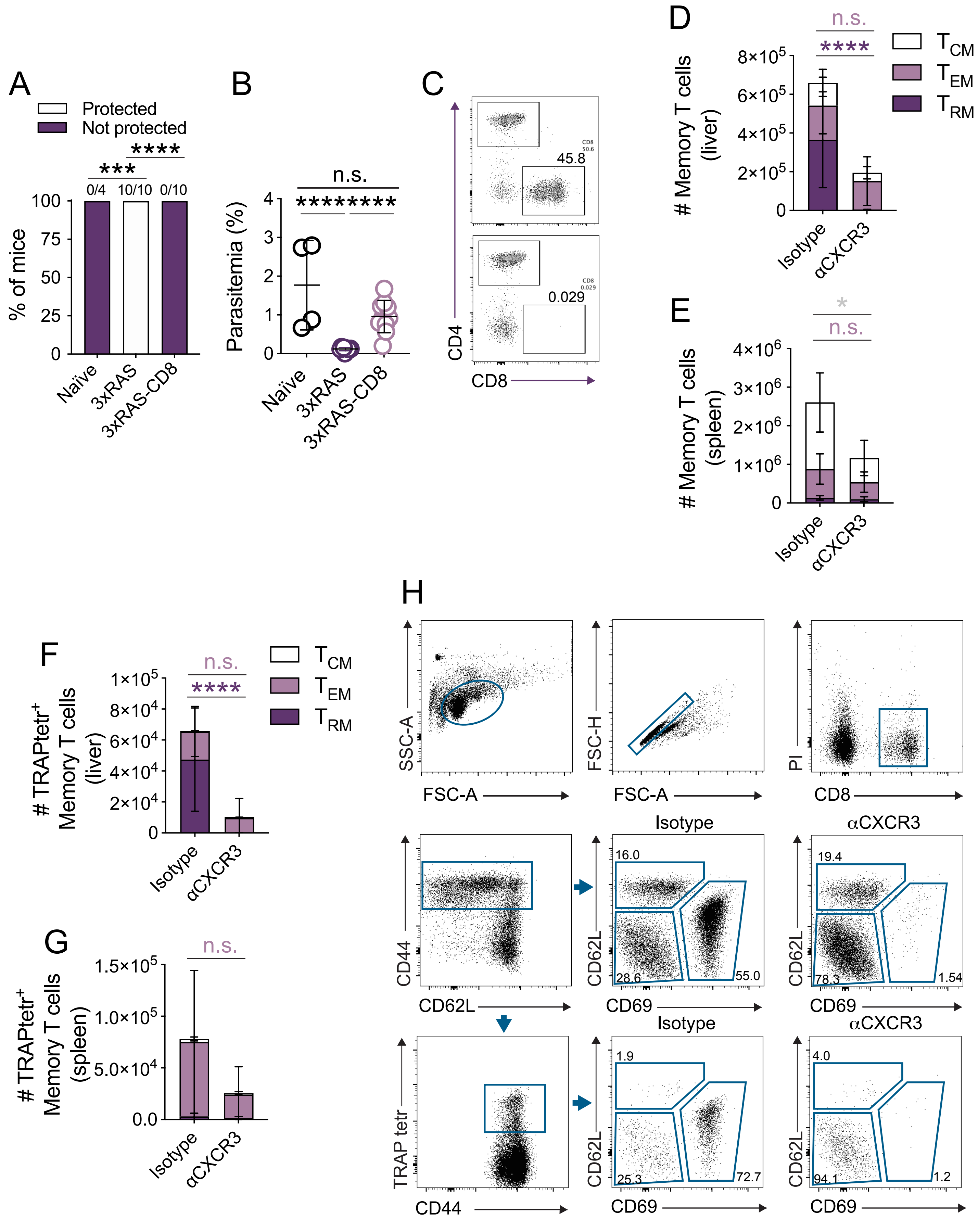

### Supplementary figure 2

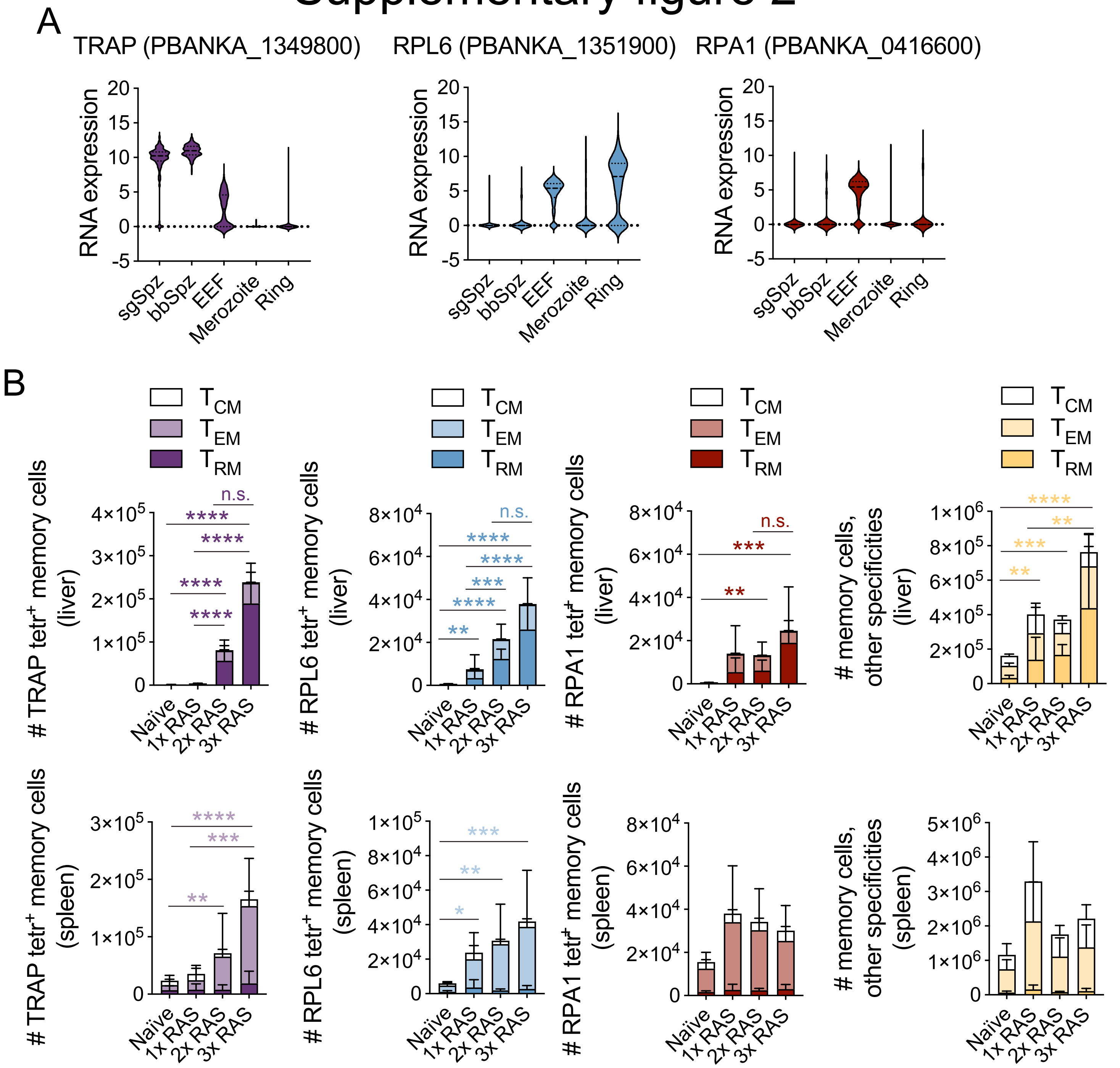

### Supplementary figure 3

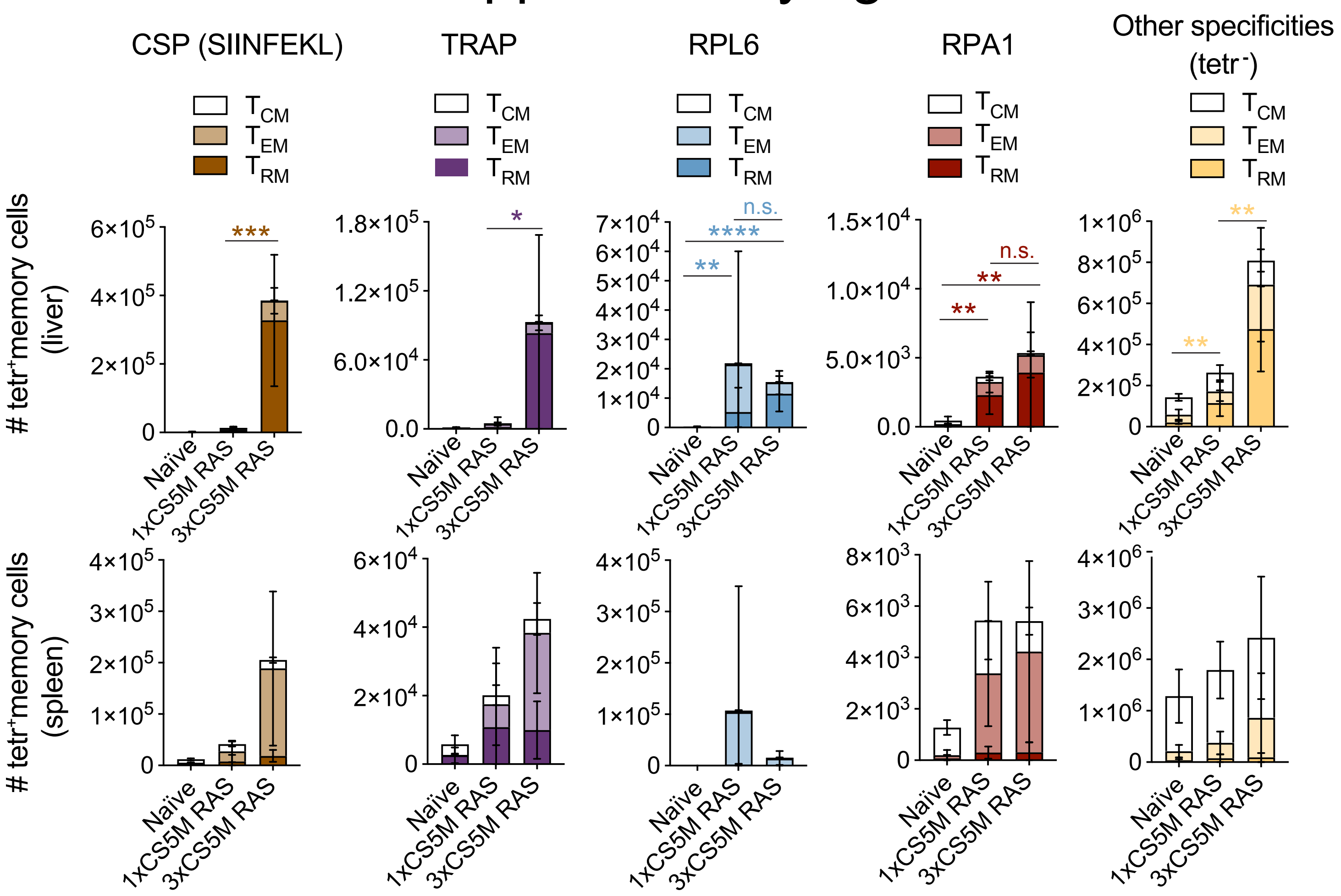


### Supplementary figure 4

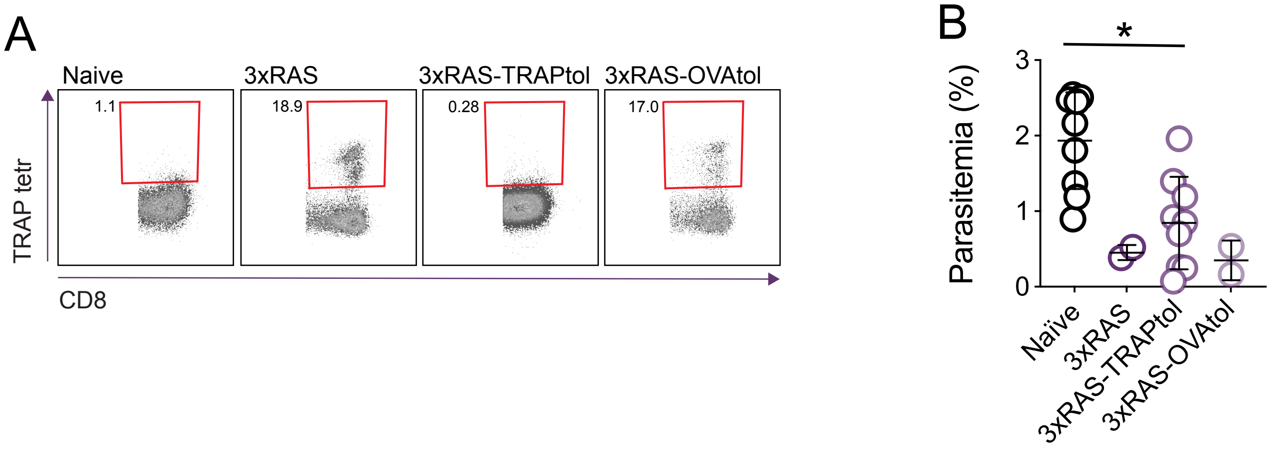

### Supplementary figure 5

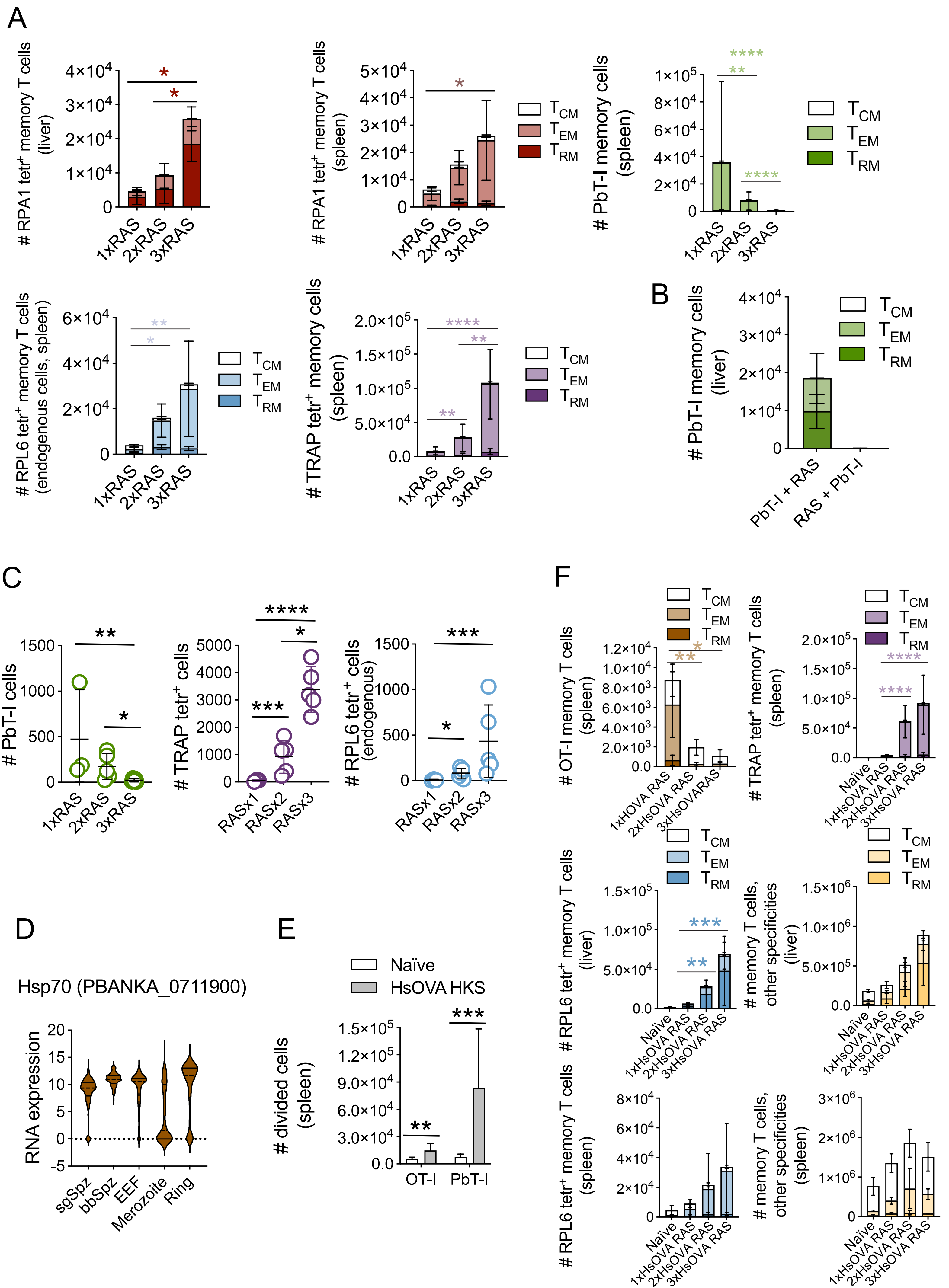

### Supplementary figure 6

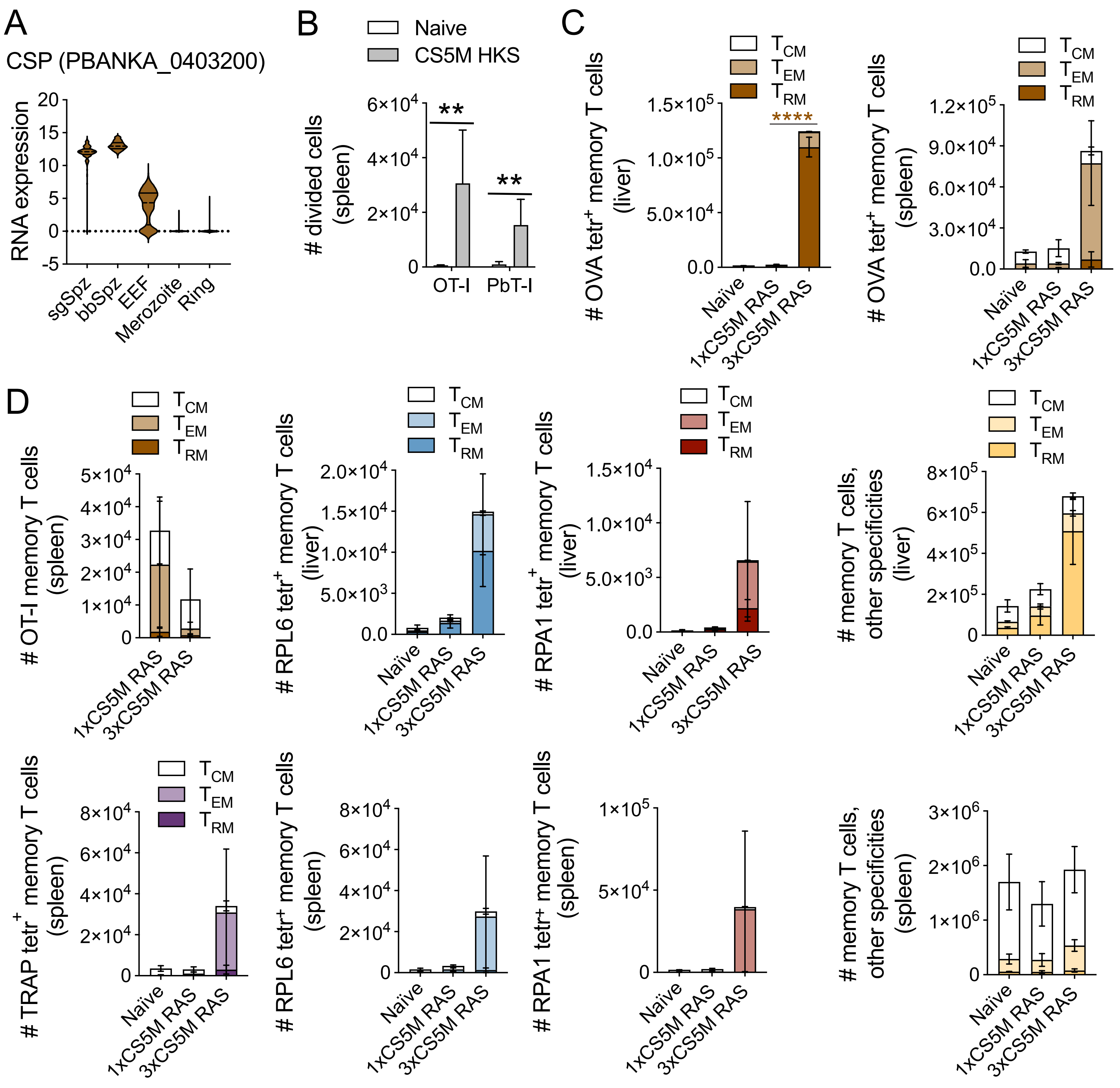

### Supplementary figure 7

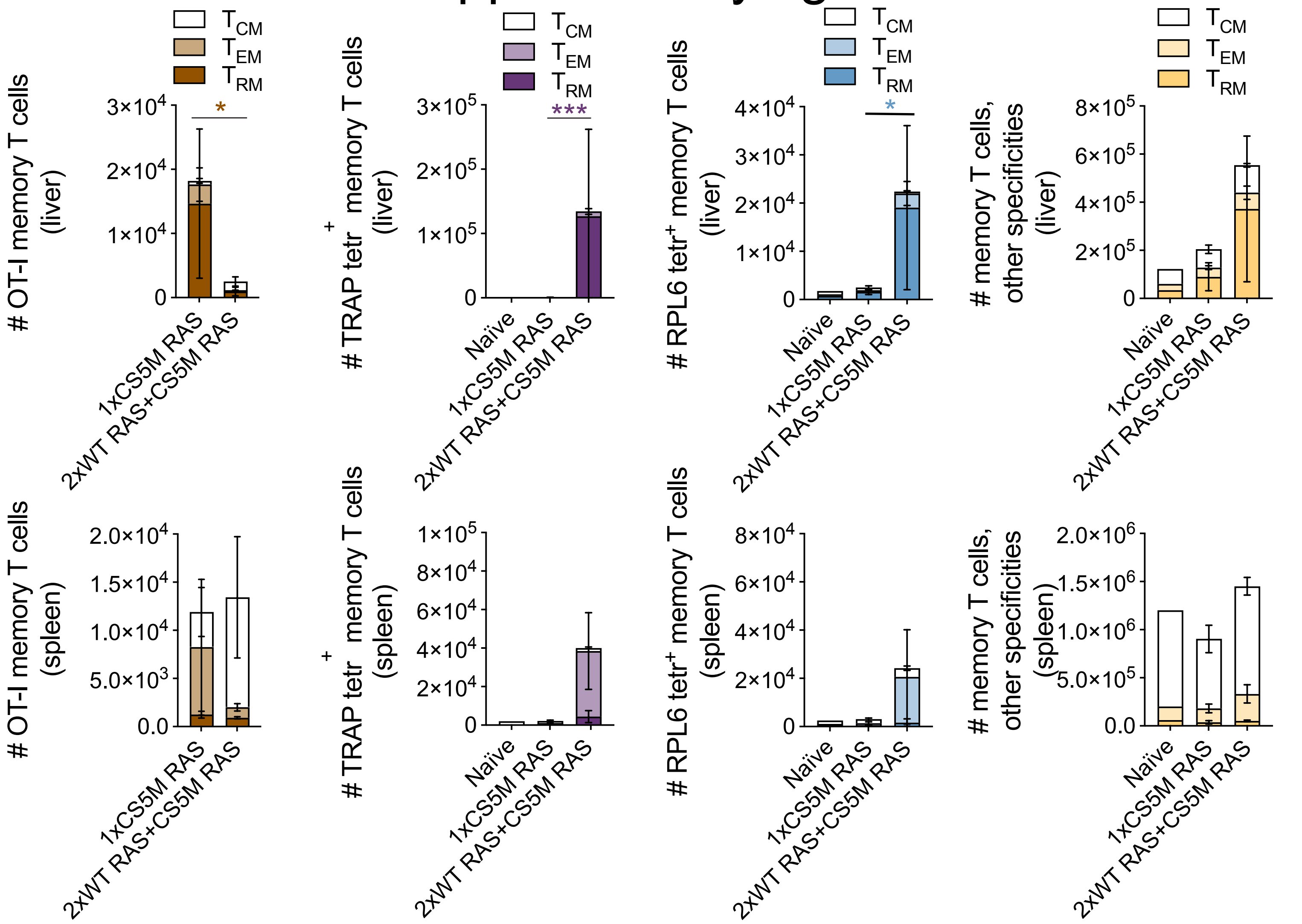


### Supplementary figure 8

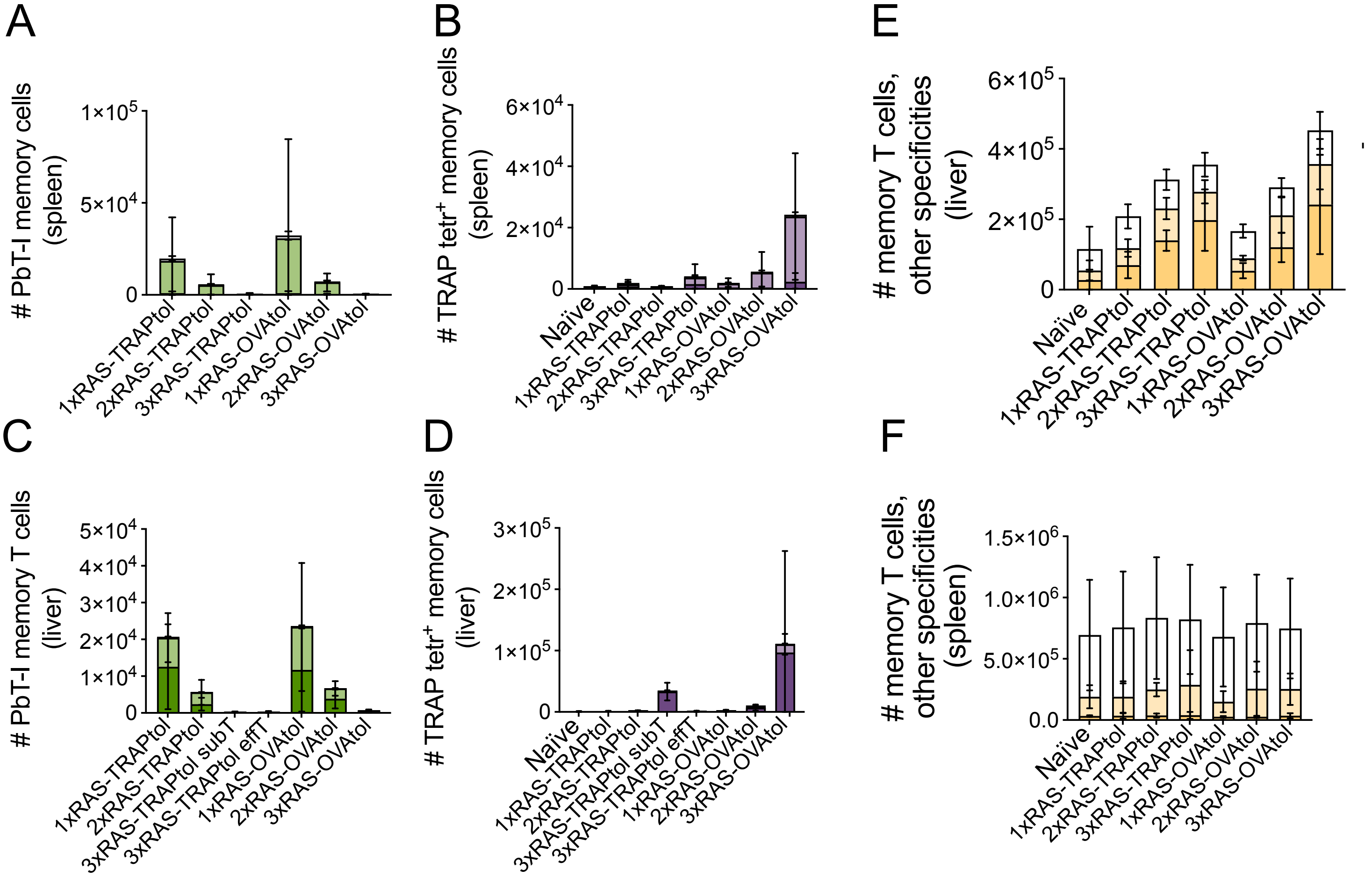


effT=efficient tolerisation of TRAP
subT=imperfect tolerisation of TRAP
